# The IRE1α-XBP1 signaling axis promotes glycolytic reprogramming in response to inflammatory stimuli

**DOI:** 10.1101/2022.10.19.512943

**Authors:** Bevin C. English, Hannah P. Savage, Scott P. Mahan, Vladimir E. Diaz-Ochoa, Briana M. Young, Basel H. Abuaita, Gautam Sule, Jason S. Knight, Mary X. O’Riordan, Andreas J. Bäumler, Renée M. Tsolis

**Author notes:** Department of Pathology, University of California San Francisco (Bevin C. English). Department of Pathobiological Sciences, School of Veterinary Medicine, Louisiana State University (Basel H. Abuaita).

## Abstract

Immune cells must be able to adjust their metabolic programs to effectively carry out their effector functions. Here, we show that the ER stress sensor IRE1α and its downstream transcription factor XBP1 enhance the upregulation of glycolysis in classically activated macrophages (CAM). The IRE1α-XBP1 signaling axis supports this glycolytic switch in macrophages when activated by LPS stimulation or infection with the intracellular bacterial pathogen *Brucella abortus*. Importantly, these different inflammatory stimuli have distinct mechanisms of IRE1α activation; while TLR4 supports glycolysis under both conditions, TLR4 is required for activation of IRE1α in response to LPS treatment but not *B. abortus* infection. Though IRE1α and XBP1 are necessary for maximal induction of glycolysis in CAM, activation of this pathway is not sufficient to increase the glycolytic rate of macrophages, indicating that the cellular context in which this pathway is activated ultimately dictates the cell’s metabolic response and that IRE1α activation may be a way to fine-tune metabolic reprogramming.

**IMPORTANCE:** The immune system must be able to tailor its response to different types of pathogens in order to eliminate them and protect the host. When confronted with bacterial pathogens, macrophages, frontline defenders in the immune system, switch to a glycolysis-driven metabolism to carry out their antibacterial functions. Here, we show that IRE1α, a sensor of ER stress, and its downstream transcription factor XBP1 support glycolysis in macrophages during infection with *Brucella abortus* or challenge with *Salmonella* LPS. Interestingly, these stimuli activate IRE1α by independent mechanisms. While the IRE1α-XBP1 signaling axis promotes the glycolytic switch, activation of this pathway is not sufficient to increase glycolysis in macrophages. This study furthers our understanding of the pathways that drive macrophage immunometabolism and highlights a new role for IRE1α and XBP1 in innate immunity.

## INTRODUCTION

It is becoming increasingly evident that the metabolism of immune cells is closely tied to their effector functions; thus, immune cells must be able to alter their metabolic programs in response to different stimuli. Macrophages have different activation states and associated metabolic programs that enable them to carry out different physiological roles. While there are likely many different activation profiles *in vivo*, one activation state that has been studied extensively are classically activated macrophages (CAM). These CAM, sometimes referred to as M1 macrophages, have a glycolysis-driven metabolism, allowing for the rapid production of ATP and antimicrobial products, such as reactive oxygen and nitrogen species (1). A variety of stimuli can induce CAMs, including certain cytokines and bacterial pathogens or products, such as LPS (1) and the intracellular pathogen *Brucella abortus* (2–4).

The endoplasmic reticulum (ER) is an organelle that plays a key role in maintaining cellular homeostasis. When ER function is perturbed, the cell experiences ER stress and initiates the unfolded protein response (UPR), a collection of linked signaling cascades, to overcome the initiating stress and return to homeostasis. The most evolutionarily conserved branch is that of the ER stress sensor IRE1α. Upon activation, IRE1α oligomerizes and trans-autophosphorylates, activating its RNase activity (5). One key function of activated IRE1α is the excision of a noncanonical intron from the unspliced XBP1 transcript (*XBP1u*), resulting in the spliced XBP1 transcript (*XBP1s*), which encodes a transcription factor that regulates a wide range of genes involved in a variety of cellular processes (6, 7).

UPR signaling is closely linked to the immune system. IRE1α signaling leads to activation of JNK (8), NF-κB (9, 10), and NOD1 and NOD2 (11, 12), while XBP1 directly regulates the expression of proinflammatory cytokines (13, 14). The UPR is activated in many different immune cells after stimulation, including T cells (15), NK cells (16), and macrophages (17–19). Many intracellular pathogens induce ER stress in their host cells (20, 21), including *Brucella* spp., which use their type IV secretion systems (T4SS) to interact extensively with the ER (22), ultimately leading to UPR activation (23–27). IRE1α has been shown to be phosphorylated upon *Brucella* infection (23, 28) and to form puncta throughout infected cells (23). Though it is well established that IRE1α plays an important role in the development and effector functions of immune cells, the links between IRE1α activation and innate immunity remain poorly understood. Thus, we set out to determine how IRE1α influences the activation of macrophages in response to inflammatory stimuli.

## RESULTS

### IRE1α supports lactate production and CAM gene expression during *B. abortus* infection or LPS stimulation in macrophages

During *in vitro* infection with *B. abortus*, macrophages shift their metabolism to be more glycolysis-driven (2–4). Consistent with this, we observed that RAW 264.7 macrophage-like cells acidify the culture media during infection, as indicated by the yellowing of the pH indicator phenol red in the media. However, we noticed that the media on IRE1α knockout (KO) RAW 264.7 cells (29) was not yellowing to the same extent as the media on wildtype (WT) cells during infection; thus, we hypothesized that the IRE1α-deficient cells were producing less lactate. Indeed, the IRE1α KO RAW 264.7 cells produce less lactate after *B. abortus* infection compared to WT cells **(Fig. 1A).** We also assessed the expression of two genes involved in glycolysis, *Glut1*, which encodes a glucose importer, and *Pfkfb3*, which encodes a glycolytic enzyme, as well as *Irg1* (also called *Acod1*), which is a marker of CAMs (30). Similar to what we observed with lactate levels, the IRE1α KO RAW cells show an impaired induction of these genes **(Fig. 1B),** suggesting that IRE1α supports the *Brucella*-induced glycolytic switch in macrophages.

**Figure 1.**
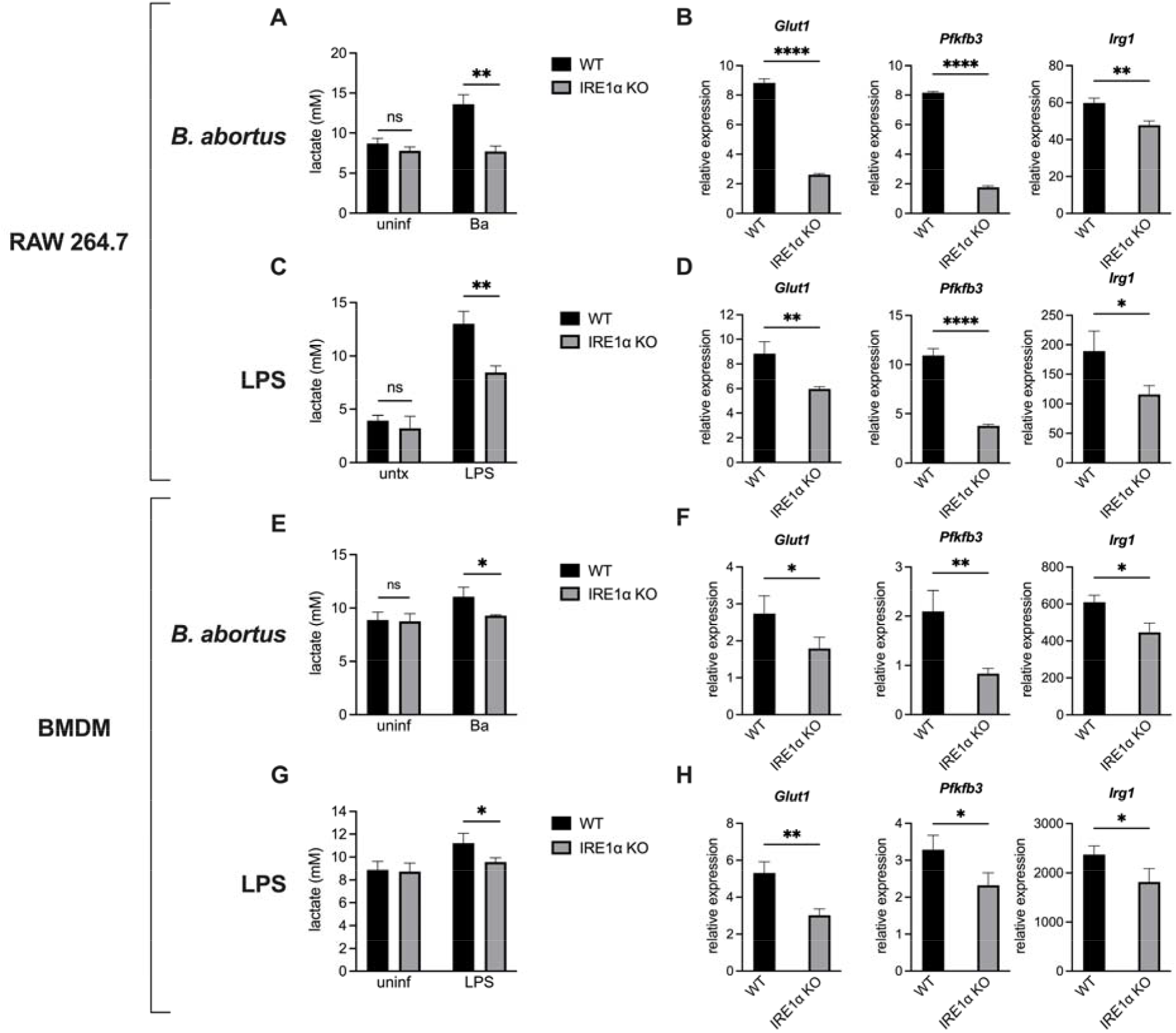
IRE1α supports lactate production and glycolytic gene expression during *B. abortus* infection or LPS stimulation in macrophages. Wildtype (WT) and IRE1α knockout (KO) RAW 264.7 cells were infected with *B. abortus* (Ba) for 48 h **(A, B)** or stimulated with 100 ng/mL *Salmonella* LPS for 24 h **(C, D).** Supernatant lactate was quantified (A, C) and relative expression of the indicated genes normalized to uninfected controls was assessed by RT-qPCR **(B, D). (E-H)** Same as panels A-D, but with WT (LysM-Cre^-^ *Ern1^fl/fl^*) or IRE1α KO (LysM-Cre^+^ *Ern1^fl/fl^*) bone marrow-derived macrophages (BMDMs). Data are presented as means of triplicate wells ± SD. * *P* ≤ 0.05, ** *P* ≤ 0.01, *** *P* ≤ 0.001, **** *P* ≤ 0.0001, ns, no statistical difference, Student’s two-tailed t-test.

It has been reported previously that IRE1α contributes to the intracellular replication of *Brucella* (23, 25, 26, 31–33), and we also observed that IRE1α-deficient macrophages had a slight reduction in bacterial burden during infection **(Fig. S1A and B).** Thus, to ensure that the reduced glycolytic shift in IRE1α KO macrophages was not secondary to reduced bacterial burden, we tested an additional stimulus. LPS is commonly used to polarize CAMs and activates IRE1α through TLR4 signaling (13, 34). When treated with LPS from *Salmonella enterica* serotype Typhimurium, a potent TLR4 agonist, IRE1α KO RAW 264.7 cells had a reduced glycolytic response **(Fig. 1C and D),** consistent with what we observed with *B. abortus* infection.

Because RAW 264.7 cells are a murine cancer-derived cell line, we wanted to confirm our findings in primary cells. To this end, we tested bone marrow-derived macrophages (BMDMs) from IRE1α conditional knockout animals (*LysM-Cre*^+/-^ *Ern1a^fl/fl^*) and their WT littermate controls (*LysM-Cre*^-/-^ *Ern1a^fl/fl^*; ref. (35)). Consistent with our observations with RAW 264.7 cells, the IRE1α-deficient BMDMs also showed a reduced lactate production and glycolytic gene expression after *B. abortus* infection or LPS treatment **(Fig. 1E-H).** Together, these data demonstrate that IRE1α supports macrophage glycolytic reprogramming in response to inflammatory stimuli.

### XBP1 supports lactate production and CAM gene expression during *B. abortus* infection or LPS stimulation in macrophages

We then wanted to determine how IRE1α was promoting glycolysis in CAMs. IRE1α is both a kinase and RNase, and we wondered which of these enzymatic functions was influencing macrophage metabolism. Treatment of macrophages with 4μ8c, which inhibits the RNase activity of IRE1α without affecting its kinase activity (36), led to reduced expression of glycolytic genes after *B. abortus* infection or LPS stimulation **(Fig. S2).** There are two major outcomes of IRE1α endonuclease activity: splicing of the unspliced XBP1 mRNA (*XBP1u*), forming the spliced XBP1 transcript (*XBP1s*), which encodes a transcription factor, and regulated IRE1α-dependent decay (RIDD), a process where specific RNA species are degraded (21). Because XBP1 regulates different metabolic states in a variety of cells (15, 16, 18, 19, 37), we chose to focus on XBP1. We used CRISPR/Cas9 to generate XBP1 KO RAW 264.7 cells **(Fig. S3).** These cells had reduced expression of *Il6*, a direct XBP1s target (13), after *Brucella* infection or LPS stimulation, further demonstrating that this pathway is activated under these inflammatory conditions **(Fig. 2B and D).** Like IRE1α KO macrophages, these XBP1 KO macrophages also had a reduced glycolytic response to *B. abortus* infection **(Fig. 2A and B)** or LPS stimulation **(Fig. 2C and D).** Thus, the IRE1α-XBP1 signaling axis promotes the glycolytic switch in macrophages in response to inflammatory stimuli.

**Figure 2.**
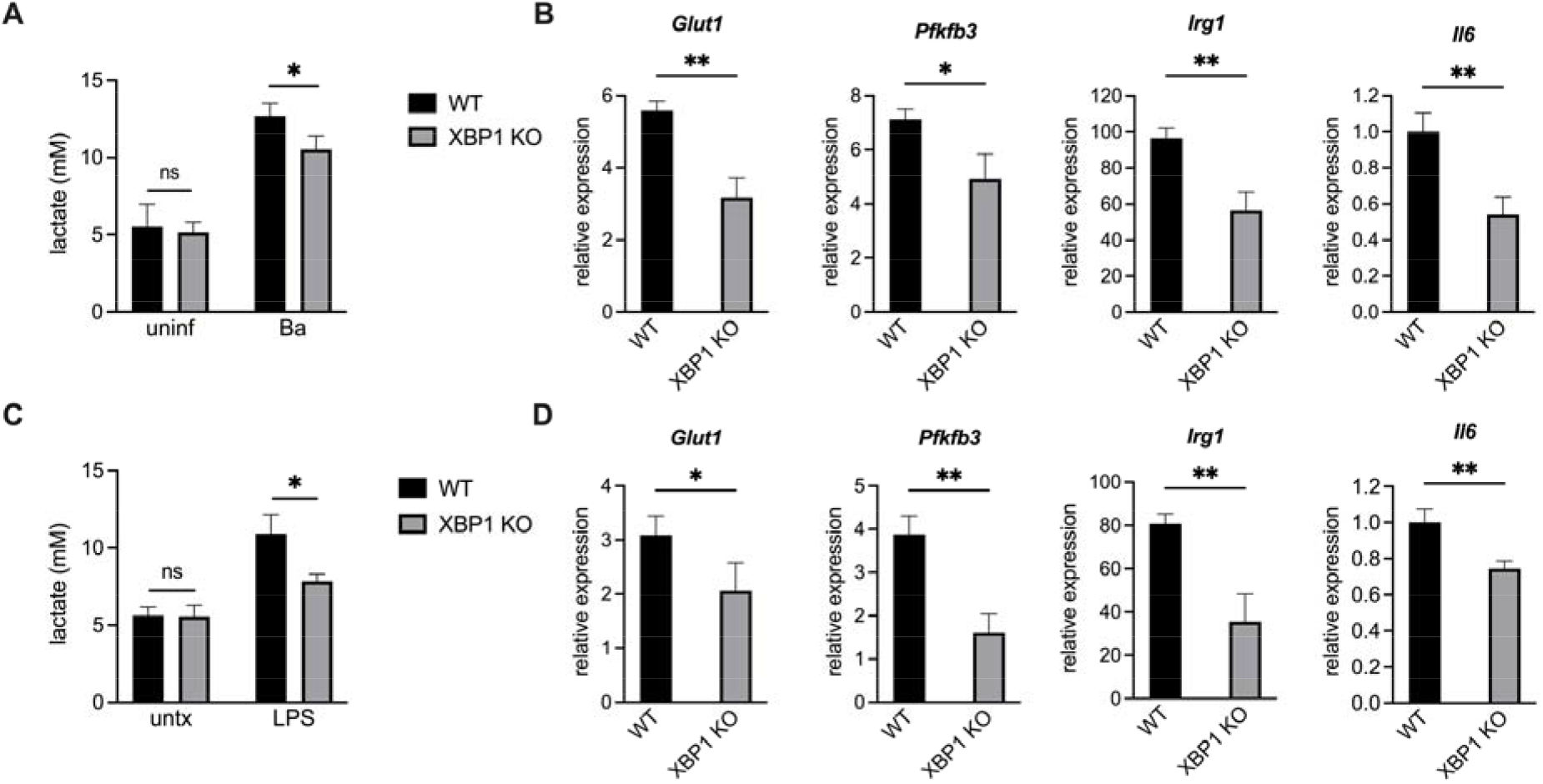
XBP1 promotes lactate production and the expression of glycolytic and inflammatory genes during *B. abortus* infection or LPS treatment. WT and XBP1 KO RAW 264.7 cells were infected with *B. abortus* (Ba) for 48 h **(A, B)** or treated with 100 ng/mL *Salmonella* LPS for 24 h **(C, D).** Supernatant lactate was quantified **(A, C)** and relative expression of the indicated genes was assessed by RT-qPCR **(B, D).** Expression levels of *Glut1, Pfkfb3*, and *Irg1* were normalized to uninfected controls. Because it is not detected in unstimulated cells, *IL6* expression was normalized to the infected or LPS-stimulated WT cells. Data are presented as means of triplicate wells ± SD. * *P* ≤ 0.05, ** *P* ≤ 0.01, ns, no statistical difference, Student’s two-tailed t-test.

### Glucose import of infected macrophages correlates with bacterial burden and is reduced in IRE1α or XBP1 KO macrophages

While infection leads to increased lactate and expression of glycolytic genes (Fig. 1–2), the magnitude of this increase was small in some cases, leading us to hypothesize that uninfected cells in our bulk assays such as qRT-PCR and lactate measurements may be masking the specific effect of *B. abortus* on the metabolic state of infected cells. To look at glycolysis on a single-cell level, we used 2-[*N*-(7-nitrobenz-2-oxa-1,3-diazol-4-yl) amino]-2-deoxy-d-glucose (2-NBDG), an unmetabolizable fluorescent glucose analog that accumulates inside cells proportionately to their glucose import rate and thus can be used to assess their glycolytic rate (38). To assess the bacterial burden of individual cells, we used a WT *B. abortus* strain that expresses mCherry (4). We observed that the glucose import rate of cells correlated with bacterial burden **(Fig. 3A and** B), suggesting that increased glycolysis is a cell-intrinsic effect of infection. Indeed, the glucose import of mock infected cells was equivalent to that of uninfected bystander cells **(Fig. 3B).** These data suggest that *B. abortus* acts directly on infected cells to promote glycolysis and that increased glycolysis is not simply a result of paracrine signaling from infected cells.

**Figure 3.**
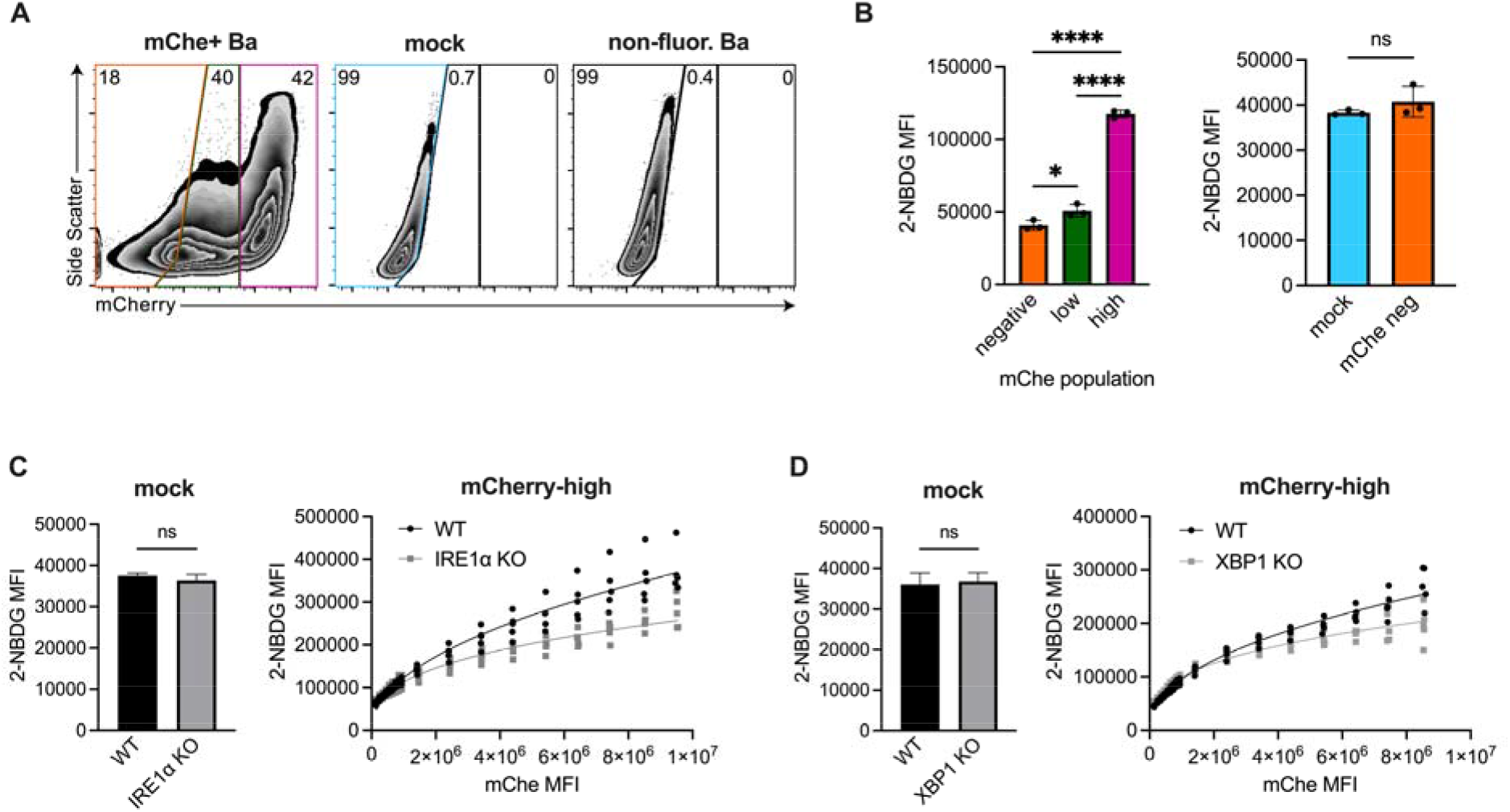
Glucose import of infected macrophages correlates with bacterial burden and is reduced in IRE1α or XBP1 KO macrophages. (A, B) WT RAW 264.7 cells were mock infected or infected with mCherry (mChe)-expressing *B. abortus* for 48 h and then stained with fluorescent glucose analog 2-NBDG. Cells were then gated based on mCherry signal. (A) Representative FACS plots showing mock or infected RAW 264.7 cells, gated on all live cells. RAW 264.7 cells infected with a wildtype non-mCherry-expressing strain is shown as an mCherry-negative control. (B) The mean fluorescence intensity (MFI) of 2-NBDG was calculated within the indicated populations. (C,D) RAW 264.7 cells of the indicated genotypes were mock infected or infected with mCherry-expressing *B. abortus* for 48 h before 2-NBDG staining. (Left) 2-NBDG MFIs of mock infected cells. (Right) 2-NBDG MFI for the mCherry-high populations after binning based on mCherry signal. Dots represent individual wells, columns are means, and error bars are SD. * P ≤ 0.05, ** *P* ≤ 0.01, *** *P* ≤ 0.001, **** *P* ≤ 0.0001, ns, no statistical difference by Student’s two-tailed t-test, except for the left panel of (B), which was analyzed by one-way ANOVA with Tukey’s post-hoc test.

We hypothesized that IRE1α-XBP1 signaling contributed to glucose import during *B. abortus* infection, as this signaling pathway was necessary for maximal expression of the glucose importer *Glut1* (**Fig. 1, 2).** Consistent with our previous data, mock-infected IRE1α and XBP1 KO macrophages had comparable glucose import compared to wildtype cells (**Fig 3C and D).** When assessing the glucose import of highly infected cells, we wanted to ensure we were comparing cells with comparable bacterial burdens. Because IRE1α supports the replication of *B. abortus* (23, 25, 26, 31–33) **(Fig. S1),** we were concerned that any observed reduction in glucose import by the IRE1α knockout macrophages could be due to reduced bacterial burden. To overcome this limitation, we binned the data across the range of mCherry signal, resulting in comparable MFIs and thus comparable bacterial burdens within each bin. We then compared the 2-NBDG signal within each bin. Across the bins, the glucose import rate of WT macrophages was higher than that of the IRE1α and the XBP1 KO macrophages **(Fig. 3C and D).** Together, these data provide more evidence that IRE1α-XBP1 signaling promotes glycolysis during *B. abortus* infection.

### IRE1α and XBP1 are required for maximal glycolytic flux after LPS stimulation

Though we demonstrated that IRE1α and XBP1 contribute to lactate accumulation, glycolytic gene expression, and glucose import after macrophage stimulation, these are indirect measurements of glycolysis, and we wanted to directly measure glycolytic flux of stimulated macrophages in real time. To this end, we assessed the extracellular acidification rate (ECAR) of our IRE1α- and XBP1-deficient macrophages after LPS stimulation. However, factors other than glycolytic rate, such as mitochondrial production of CO_2_, can contribute to ECAR; thus, we also assessed the <u>p</u>roton <u>e</u>fflux <u>r</u>ate from glycolysis (glycoPER) specifically. Consistent with our previous data, the KO macrophages showed reduced ECAR and glycoPER after LPS stimulation (Fig. 4), further demonstrating that the IRE1α-XBP1 signaling axis promotes glycolytic flux.

**Figure 4.**
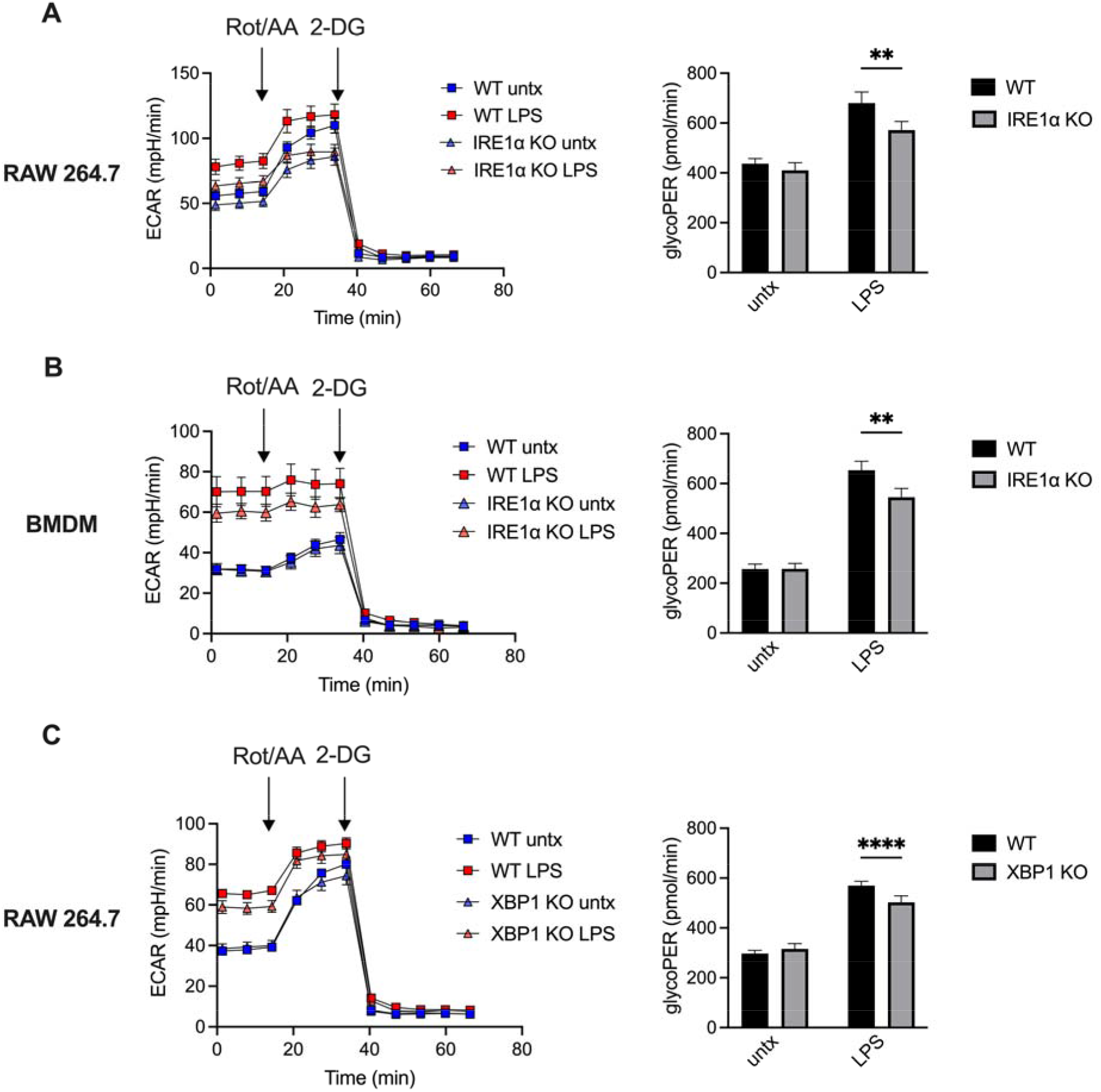
IRE1α and XBP1 support glycolysis after LPS stimulation. **(A)** WT or IRE1α KO RAW 264.7 cells, (B) WT (LysM-Cre^-^ *Ern1^fl/fl^*) or IRE1α KO (LysM-Cre^+^ *Ern1^fl/fl^*) BMDMs, and (C) WT or XBP1 KO RAW 264.7 cells were stimulated with 100 ng/mL *Salmonella* LPS for 6 hours before the assessing extracellular acidification rate (ECAR), with rotenone/antimycin A (Rot/AA) and 2-doxyglucose (2-DG) treatments as indicated (left). The proton efflux rate from glycolysis (glycoPER) was calculated as a more specific assessment of glycolytic flux (right).

### TLR4 supports glycolysis in macrophages but is not required for IRE1α activation during *B. abortus* infection

We then wondered if both *Salmonella* LPS and *B. abortus* were activating the IRE1α-XBP1 signaling pathway in the same manner. LPS activates IRE1α via TLR4 (13, 34), and *Salmonella* LPS is a strong TLR4 agonist. Though *Brucella* spp. have a modified LPS that is a weak TLR4 agonist (39, 40) and encode an effector that downregulates TLR4 signaling during infection (41, 42), TLR4 has been shown to play a role in the response to *Brucella* infection (43, 44). Thus, we generated BMDMs from WT and TLR4 KO mice. As expected, TLR4 KO BMDMs show a severely attenuated glycolytic response to LPS stimulation **(Fig. 5C and D).** After *B. abortus* infection, TLR4 KO BMDMs also show a decreased upregulation of glycolysis **(Fig. 5A and B),** which is intriguing since *B. abortus* reduces activation of TLR4 by its LPS during infection (45, 46).

**Figure 5.**
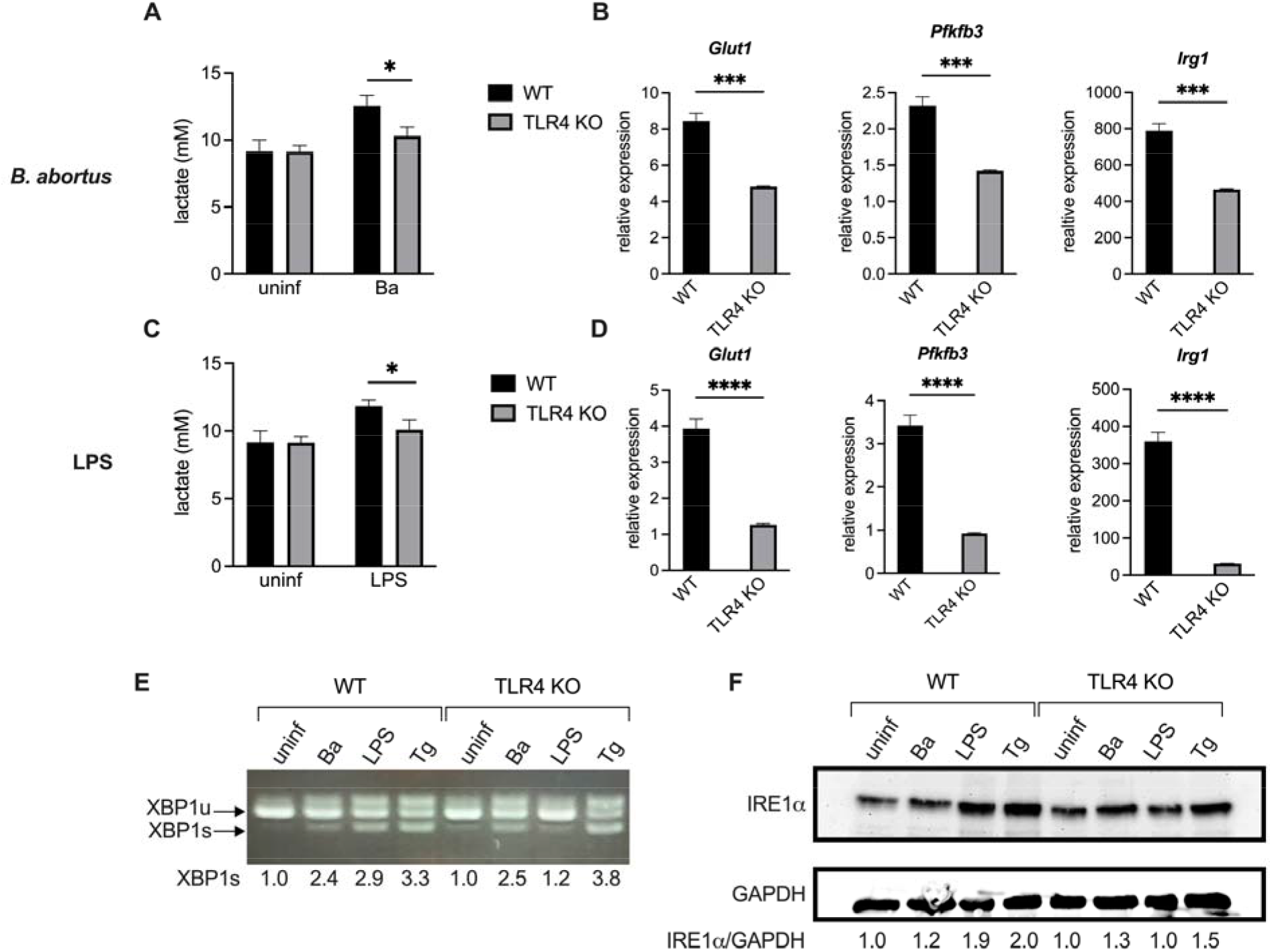
TLR4 supports glycolysis but not activation of IRE1α during *B. abortus* infection. BMDMs from WT or TLR4 KO mice were infected with *B. abortus* (Ba) for 48 h, treated with 100 ng/mL *Salmonella* LPS for 24 h, or treated with 250 nM thapsigargin (Tg) for 24 h. **(A, C)** Supernatant lactate was quantified. (B, D) Relative expression of the indicated genes normalized to uninfected controls was assessed by RT-qPCR. **(E)** *XBP1* splicing was assessed via non-quantitative RT-PCR. Densitometry of the XBP1s band relative to uninfected for each genotype is reported below. (F) IRE1α protein levels were assessed by western blot. Densitometry of the IRE1α band normalized to the GAPDH band and relative to uninfected for each genotype is reported below. Lactate and expression data are presented as means of triplicate wells ± SD. * *P* ≤ 0.05, ** *P* ≤ 0.01, *** *P* ≤ 0.001, **** *P* ≤ 0.0001, Student’s two-tailed t-test.

While these data demonstrate that TLR4 supports the induction of glycolysis in classically activated macrophages, we next wanted to determine if TLR4 was activating the IRE1α-XBP1 signaling axis or acting in a parallel pathway. We examined IRE1α activation by assessing *XBP1* splicing and IRE1α levels, since IRE1α activation leads to IRE1α upregulation in a positive feedback loop (47). As expected, LPS treatment of TLR4 KO macrophages failed to induce significant *XBP1* splicing or IRE1α upregulation. However, TLR4 KO macrophages showed robust *XBP1* splicing and modest IRE1α upregulation during *B. abortus* infection **(Fig. 5E and F),** demonstrating that TLR4 is not required for IRE1α activation during *B. abortus* infection.

### Maximal glucose import by macrophages is dependent on the type IV secretion system during *B. abortus* infection

Because TLR4 is not required for the IRE1α-mediated induction of glycolysis during *B. abortus* infection, we then interrogated how *B. abortus* was promoting glycolysis in macrophages. *B. abortus* uses its type IV secretion system (T4SS) to interact extensively with the host cell ER, resulting in robust intracellular replication and the induction of ER stress and subsequent IRE1α activation (22). Thus, we investigated the role of the T4SS in *Brucella*-induced glycolytic induction in macrophages. We expressed mCherry in our *virB2* mutant bacteria, which lack the T4SS, and in the complemented strain (48). Because the T4SS is required for intracellular replication, we increased the multiplicity of infection (MOI) used for the T4SS mutant, as we have previously observed that this increased MOI results in macrophages containing a high burden of the T4SS mutant (49).

Even though all strains express the same level of fluorescence **(Fig. S4A),** the mCherry signal of cells infected with the T4SS mutant at a high MOI did not match that of macrophages infected with the complemented strain (Fig. S4B), which was equivalent to that of wildtype **(Fig. S4C).** For the mCherry-high population of RAW cells, the mCherry mean fluorescence intensity (MFI) was significantly higher for the macrophages infected with the complemented strain **(Fig. S4B).** This suggests that increasing the MOI cannot fully compensate for the intracellular replication defect of the T4SS mutant. Thus, 2-NBDG uptake by all cells highly infected with either the mutant or complemented strain could not be compared, since 2-NBDG uptake is correlated with the level of infection **(Fig. 3B).** To overcome this difference, we compared cells infected with either the mutant or the complemented strain in two ways. First, we compared glucose uptake among mCherry-low cells. For the mCherry-low cells, the mCherry mean fluorescence intensity (MFI) was significantly higher for the cells infected with the *virB2* mutant than those infected with the complemented strain. However, despite this disparity in bacterial burdens, the macrophages showed equivalent glucose uptake, suggesting that the T4SS promotes the glycolytic switch in infected macrophages **(Fig. 6A).** Next, for the more highly infected cells, we again binned the data across the range of mCherry signal. Consistent with our previous observation, as bacterial burden increased, the 2-NBDG signal increased. And for all bins, the glucose import rate of macrophages infected with the complemented strain was consistently higher than that of macrophages infected with the T4SS mutant **(Fig. 6B).** Together, these data demonstrate that the T4SS contributes to the glycolytic switch of infected macrophages.

**Figure 6.**
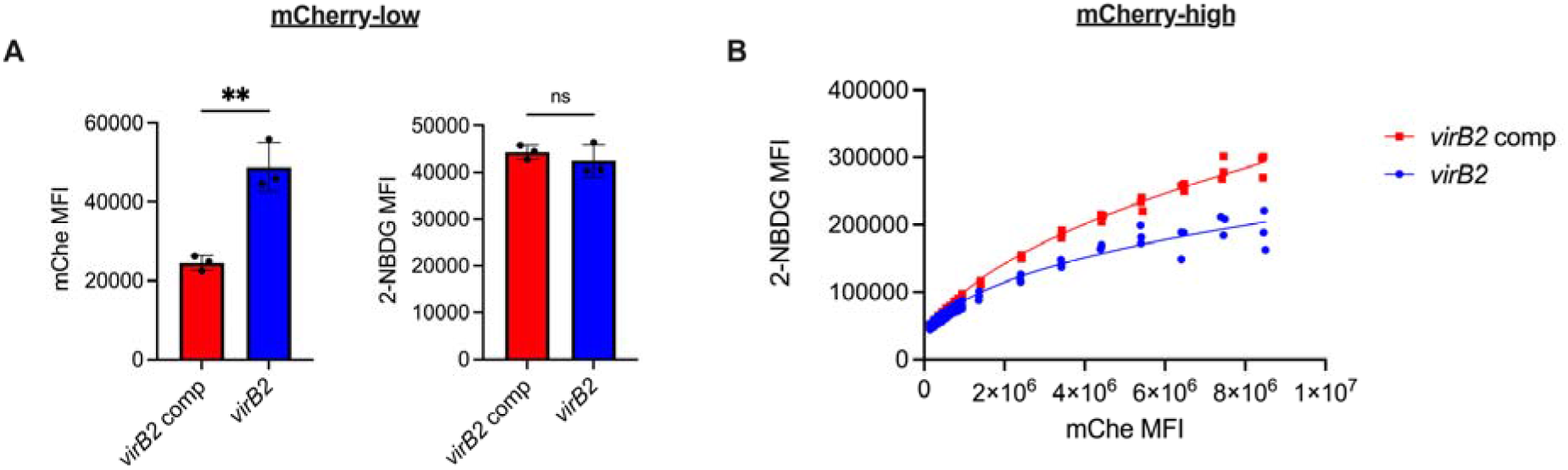
The type IV secretion system promotes glucose import by macrophages during *B. abortus* infection. RAW 264.7 cells were infected with the T4SS-deficient mCherry-expressing *virB2* mutant at an MOI of 2000 or the mCherry-expressing complemented *virB2* strain at an MOI of 100 and then stained with 2-NBDG after 48 h. (A) MFIs of mCherry and 2-NBDG for the mCherry-low population of RAW 264.7 cells infected with the indicated *B. abortus* strains. (B) 2-NBDG MFI for the mCherry-high populations infected with the indicated *B. abortus* strains after binning based on mCherry signal. Dots represent individual wells, columns are means, and error bars are SD. ** *P* ≤ 0.01, ns, no statistical difference by Student’s two-tailed t-test.

### Impaired induction of glycolysis does not necessarily impact intracellular replication of *B. abortus*

We and others have shown that IRE1α contributes to the intracellular replication of *Brucella* ((23, 25, 26, 31–33); **Fig S1).** Intriguingly, it has also been shown that glycolysis in the infected macrophage and lactate catabolism by *Brucella* also support intracellular replication (2). We wondered if IRE1α was promoting intracellular replication by increasing glycolysis. *B. abortus* showed no replication defect in XBP1 or TLR4 KO macrophages **(Fig. S5A, B),** despite those cells’ reduced glycolytic induction. Thus, reducing the glycolytic rate of infected cells does not necessarily impair the intracellular growth of *B. abortus*, suggesting that IRE1α promotes the intracellular replication of *B. abortus* independently of its role in the glycolytic switch.

### Activation of the IRE1α-XBP1s pathway is not sufficient to increase glycolysis

Having established that the IRE1α-XBP1 signaling axis promotes glycolysis in CAM, we then investigated if activation of this pathway was sufficient to cause increased glycolysis. We treated RAW 264.7 cells with the chemical ER stress inducers tunicamycin and thapsigargin, which led to robust *XBP1* splicing **(Fig. 7A).** However, neither of these treatments led to an upregulation of glycolytic genes **(Fig. 7B),** suggesting that IRE1α activation is not sufficient to increase glycolysis.

**Figure 7.**
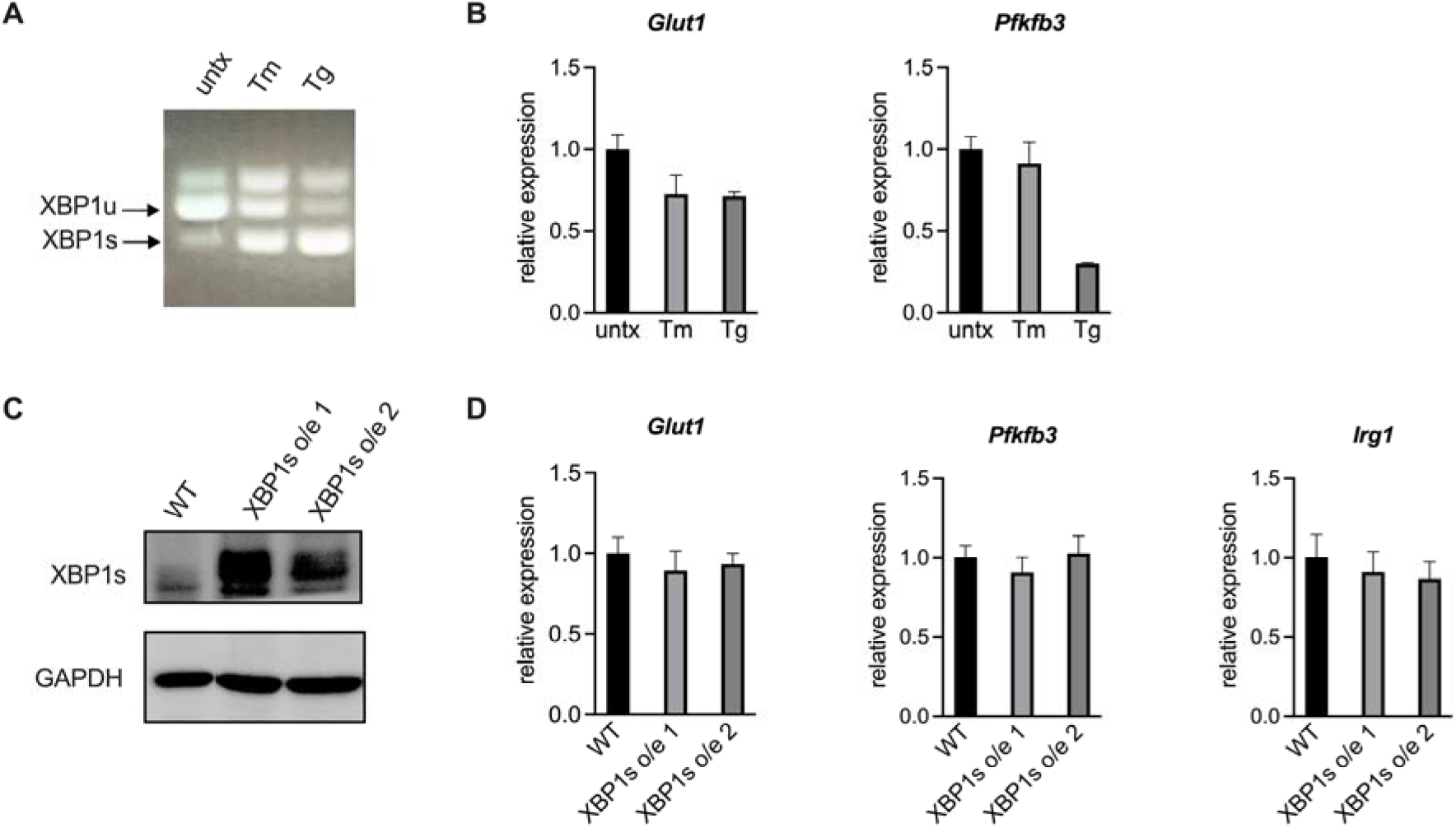
Activation of the IRE1α-XBP1 signaling axis is not sufficient to increase glycolysis in macrophages. **(A, B)** RAW 264.7 cells were treated with 200 ng/mL tunicamycin (Tm) or 50 nM thapsigargin (Tg) for 24 h. XBP1 splicing was assessed by non-quantitative RT-PCR (A), and expression of the indicated genes normalized to untreated controls was assessed by RT-qPCR **(B). (C, D)** Two independent XBP1s overexpression (o/e) RAW 264.7 cell lines were generated. XBP1s protein levels were assessed by western blotting (C), and expression of the indicated genes normalized to wildtype RAW 264.7 was assessed by RT-qPCR (D). Data are means of triplicate wells ±SD.

Tunicamycin and thapsigargin are potent ER stress inducers that activate all three branches of the UPR. To look more specifically at the IRE1α-XBP1 signaling axis, we overexpressed XBP1s in two independently generated RAW 264.7 cell lines **(Fig. 7C).** As we observed with chemical IRE1α activation, overexpression of XBP1s was not sufficient to upregulate glycolytic genes or the CAM marker *Irg1* **(Fig. 7D).** Together, these data demonstrate that activation of the IRE1α-XBP1s signaling pathway is not sufficient to increase glycolysis in macrophages.

## DISCUSSION

A cell must utilize the right metabolic pathways to optimally perform its effector functions, and the ability to modulate metabolic processes is essential for cells that must respond to different stimuli, especially immune cells. Here, we show that the ER stress sensor IRE1α and its downstream regulator XBP1s contribute to metabolic reprogramming of macrophages by promoting glycolysis in response to inflammatory stimuli. This occurs when IRE1α is activated in a TLR4-dependent manner, such as with LPS, or a TLR4-independent manner, such as with *B. abortus* **(Fig. S6).** While TLR4 is not required for IRE1α activation during *B. abortus* infection, TLR4-deficient macrophages show a reduction in glycolytic flux during infection, suggesting that TLR4 can support glycolysis via IRE1α-dependent and IRE1α-independent mechanisms. TLR4 signaling has been shown to lead to the accumulation of HIF-1α, a key transcriptional regulator of CAM (50), and the IRE1α-XBP1 signaling axis enhances HIF-1α transcriptional activity without affecting HIF-1α protein levels in cancer cells (37).

In this study, we chose to focus on the IRE1α-XBP1 signaling axis; however, because IRE1α activation has effects other than XBP1 splicing, we cannot rule out a role for these other IRE1α functions in metabolic reprogramming. Indeed, while IRE1α-deficient cells showed no increase in lactate production after LPS stimulation or *Brucella* infection, the XBP1-deficient cells produced an intermediate level of lactate, suggesting that IRE1α may also promote glycolysis via XBP1-independent mechanisms **(Fig. 2A, C)** For example, IRE1α phosphorylation leads to JNK activation (8), and JNK signaling promotes the Warburg effect in cancer cells (51). In addition to splicing the *XBP1* transcript, activated IRE1α also degrades specific RNA species in a process called regulated IRE1α-dependent decay, or RIDD. Intriguingly, RIDD, which contributes to the intracellular survival of *Brucella* (28), influences the metabolism of cancer cells (52) and thus may also be affecting the metabolism of CAM.

While our results demonstrate that the IRE1α-XBP1 signaling axis supports glycolysis in CAM, it is unclear if this pathway is also involved in other metabolic changes during macrophage activation. *Brucella* infection leads to the production of mitochondrial ROS (53, 54), mitochondrial fragmentation (55), and decreased mitochondrial metabolism (2), while IRE1α activation leads to increased mitochondrial ROS during infection with an attenuated *B. abortus* strain (53) or multidrug-resistant *Staphylococcus aureus* (29). However, during *B. abortus* infection, ROS production and subsequent IL-1β production are XBP1-independent (53), but here we show that XBP1 contributes to glycolysis. On the other hand, XBP1 inhibits mitochondrial function in tumor-associated T cells (15). Future studies will investigate how the IRE1α-XBP1 signaling axis affects mitochondrial function during CAM polarization.

One interesting aspect of this study is the observation that reduced glycolytic induction in macrophages is not sufficient to alter *B. abortus* intracellular replication. While IRE1α, XBP1, and TLR4 KO macrophages all showed reduced glycolysis during infection, only the IRE1α-deficient macrophages showed a reduced bacterial burden, suggesting that IRE1α supports *B. abortus* replication independently of enhanced glycolysis. *Irg1* has been implicated in the control of *Brucella in vivo* (56), but we did not observe enhanced replication in macrophages with reduced *Irg1* expression. It has also been reported that inhibition of host cell glycolysis and lactate production with potent small molecule inhibitors impairs *Brucella* replication (2). However, it is worth noting that IREα and XBP1 KO macrophages are still upregulating glycolytic genes **(Fig. 1B, D, F, H; Fig. 2B, D),** glucose import **(Fig. 3C, D),** and glycolytic flux **(Fig. 4)** under inflammatory stimuli, just to a lesser extent than WT cells. Lactate utilization is required for robust intracellular replication during *in vitro* infection (18). However, during chronic infection *in vivo, B. abortus* favors alternatively activated macrophages (AAM), which have markedly different metabolism compared to CAM, due to increased glucose availability (4). We believe the ability *of B. abortus* to replicate in cells with different metabolic states contributes to its success as a pathogen. Indeed, different metabolic states of the host cell contribute to the replication of other intracellular pathogens, including *Chlamydia trachomatis* (57), *Salmonella enterica* (58), and *Legionella pneumophila* (59).

It is clear the ER plays a central role in both sensing and directing different metabolic processes and that IRE1α activation can have profound effects on cellular metabolism (60). However, those effects are very context dependent. In NK cells, IRE1α-XBP1 signaling during viral infection drives oxidative phosphorylation mediated by c-Myc (16), while XBP1s inhibits mitochondrial function in tumor-infiltrating T cells (15). In breast cancer cells, XBP1s cooperates with HIF-1α to directly regulate many glycolytic genes, including the glucose importer *Glut1* (37). In obese mice, the IRE1α-XBP1 axis represses AAM polarization (18), and in non-obese mice, it contributes to the mixed phenotype of tumor-associated macrophages, regulating the expression of both CAM and AAM markers (19). By demonstrating that IRE1α-XBP1 signaling is required for robust glycolytic induction in macrophages in response to different inflammatory stimuli, we have provided another context in which this critical signaling pathway plays an important role in immunometabolism.

## MATERIALS AND METHODS

### Bacterial strains and culture conditions

Bacterial strains in this study are the virulent wildtype *Brucella abortus* 2308; its isogenic mCherry+ strain MX2 (4); the T4SS-deficient *virB2* mutant ADH3 (48); its isogenic mCherry+ BCE4; the *virB2* complemented strain ADH8 (48); and its isogenic mCherry÷ BCE5. MX2, BCE4, and BCE5 each have an insertion of the pKSoriT-*bla-kan-PsojA-mCherry* plasmid (61). BCE4 and BCE5 were generated via conjugation with S17 *E. coli* bearing the mCherry plasmid; clones that were kanamycin resistant and fluorescent were selected and the insertion site was validated by multiplex PCR (primers listed in table S1.)

All *B. abortus* strains were cultured on blood agar plates (UC Davis Veterinary Medicine Biological Media Services) for 3 days at 37°C with 5% CO_2_. *B. abortus* was then cultured overnight at 37°C with aeration in tryptic soy broth (TSB; BD Difco), then subcultured in acidic EGY (pH 5.5) for 4 hours at 37°C with aeration prior to macrophage infections. To confirm equivalent fluorescent signals between MX2, BCE4, and BCE5, each strain was grown in triplicate overnight cultures in TSB, then mCherry fluorescence was measured on a GloMax Explorer Microplate Reader (Promega) and viable cells enumerated by colony forming units (CFU) after plating on tryptic soy agar (BD Difco) plates and incubating at 37°C with 5% CO_2_ for 3 days. All work with *B. abortus* was performed at biosafety level 3 and was approved by the Institutional Biosafety Committee at the University of California, Davis.

### Mammalian cell culture

RAW 264.7 murine macrophage-like cells (TIB-71; ATCC) and their derivatives were cultured in RPMI 1640 media (Gibco) supplemented with 10% heat-inactivated fetal bovine serum (FBS) at 37°C, 5% CO_2_. Bone marrow-derived macrophages (BMDMs) were generated as previously described (4). Briefly, bone marrow cells from femurs and tibiae from 6 to 8 week old female C57BL/6J (Jackson Labs, stock 000664), TLR4 KO (Jackson Labs, stock 029015), LysM-Cre^+^ *Ern1^fl/fl^*, and LysM-Cre^-^ *Ern1^fl/fl^* (ref. (35)) mice were isolated and maintained in RPMI 1640 supplemented with 10% FBS (Gibco), 30% L929 cell supernatant, and GlutaMAX™ (Gibco) at 37°C, 5% CO_2_ for 7 days before use in *in vitro* assays. The same media was used for all subsequent BMDM experiments. Lipopolysaccharides (LPS) from *Salmonella enterica* serotype Typhimurium (Sigma) and tunicamycin (Sigma) were reconstituted in D-PBS (Gibco). Thapsigargin (Sigma) and 4μ8c (Sigma) were reconstituted in DMSO. 2-NBDG ((2-(*N*-(7-Nitrobenz-2-oxa-1,3-diazol-4-yl)Amino)-2-Deoxyglucose) (Thermo) was reconstituted in 100% ethanol.

### Generation of XBP1 knockout and overexpression cell lines

To generate XBP1 knockout (KO) cells, complementary oligos (table S1) forming the non-targeting control (NTC) sgRNA (5’ TCCTGCGCGATGACCGTCGG 3’) and the XBP1-targeting sgRNA (5’ CGGCCTTGTGGTTGAGAACC 3’) were phosphorylated, annealed, and ligated into BsmBI-digested pXPR_001 (62), resulting in pBCE44 and pBCE41, respectively. To generate lentiviral particles, pBCE44 or pBCE41 were co-transfected with psPAX2 (Addgene, plasmid 12260) and pMD2.G (Addgene, plasmid 12259) using Xfect (Takara Bio) into HEK-293T cells (CRL-2316, ATCC) grown in DMEM (Gibco) with 10% FBS (Gibco). Lentiviral particles were concentrated from clarified supernatants using Lenti-X Concentrator (Takara Bio). RAW 264.7 cells were transduced with the lentiviral particles and then selected with 4 μg/mL puromycin (Gibco). After 5 days of selection, single cells were plated by serial dilution in 96-well plates for clonal selection and gDNA was extracted from the remaining pool using the DNeasy kit (Qiagen). The *XBP1* locus was amplified by PCR and sequenced by Sanger sequencing using the primers listed in table S1, and cutting efficiency was estimated by TIDE analysis (63). After confirming efficient disruption, gDNA was prepared from clonal lines using QuickExtract (Lucigen), and the *XBP1* locus was amplified by PCR and sequenced by Sanger sequencing. Putative knockouts were further validated by western blot analysis and by measuring expression of *ERdJ4*, an XBP1s-specific target, after thapsigargin treatment.

To generate XBP1 overexpression (o/e) lines, *XBP1s* was amplified from cDNA generated from RAW 264.7 cells treated with thapsigargin and cloned into a modified pENTR1A ((64); Addgene, plasmid 17398) using Gibson Assembly Master Mix (New England BioLabs). After sequence validation by Sanger sequencing, *XBP1s* was cloned into the mammalian expression vector pLENTI CMV Puro Dest (Addgene, plasmid 17452) using LR Clonase II (Invitrogen). Lentiviral particles were generated and RAW 264.7 cells were transduced as described above such that two separate XBP1s o/e lines were generated. XBP1s overexpression was confirmed by western blot analysis.

### Macrophage infections

For most infections, one day prior to infection, RAW 264.7 cells were seeded in 24-well plates at 5 x 10^4^ cells per well and BMDMs were seeded at 1.5 x 10^5^ cells per well in 24-well tissue culture plates. For flow cytometry experiments, RAW 264.7 cells were seeded at 2 x 10^5^ cells per well in 6-well tissue culture plates. For infection of RAW 264.7 cells, *B. abortus* was opsonized for 30 minutes at room temperature with 20% antiserum in PBS++ prepared from male C57BL/6J mice infected with *B. abortus* 2308 for 2 weeks. For inoculum preparation, the bacteria were washed in D-PBS (Gibco), diluted in the appropriate cell culture media, and added to the macrophages at multiplicity of infection (MOI) of 100 unless otherwise indicated. The tissue culture plates were then centrifuged at 210 x *g* for 5 minutes to synchronize infection. After a phagocytosis period of 30 minutes at 37°C in 5% CO_2_, the cells were washed twice with D-PBS and then incubated with 50 μg/mL gentamycin (Gibco) in the appropriate culture media for 30 minutes at 37°C in 5% CO_2_, after which the media was replaced with gentamycin-free media. To examine intracellular replication by CFUs, infected macrophages were lysed in 0.5% Tween-20 at the indicated time points. The lysates were serially diluted in D-PBS and spread on TSA plates, which were then incubated at 37°C and 5% CO_2_ for 3-5 days before colony enumeration. For lactate quantification, cell culture supernatants were sterile-filtered through 0.22 μm filters and stored at −80°C until use. Lactate levels were measured using the Lactate Colorimetric Assay Kit II (BioVision) according to the manufacturer’s protocol.

### 2-NBDG assay

RAW 264.7 cells of the indicated genotypes were infected as described above at an MOI of 100 for 2308, ADH8, MX2, and BCE5 or at an MOI of 2000 for ADH3 and BCE4. After 48 hours, cells were washed three times with D-PBS and collected by scraping, Viable cells were counted on a hemacytometer using trypan blue. One million viable cells were incubated with 300 nM 2-NBDG in glucose-free DMEM (Gibco) with 10% FBS for 45 minutes, then stained with LIVE/DEAD Fixable Aqua (ThermoFisher) in D-PBS for 15 minutes, then fixed in CytoFix (BD Biosciences) for 30 minutes. Cells were then run on a CytoFLEX Flow Cytometer (Beckman Coulter) and data were analyzed using FlowJo (10.8.0). Non-fluorescent *B. abortus* strains were used to inform gating strategies. When indicated for the mCherry-high cells, the mCherry signal was binned by equal units within each log across the population (e.g., 1 x 10^6^, 2 x 10^6^, 3 x 10^6^, etc.).

### Seahorse analysis

2 x 10^4^ RAW 264.7 cells or 3 x 10^4^ BMDMs were seeded in Seahorse XF96 Cell Culture Microplates. The next day, the cells were treated with 100 ng/mL LPS for 6 hours in the appropriate media. The cells were assessed on a Seahorse XFe96 Analyzer (Agilent) using the Seahorse XF Glycolytic Rate Assay Kit according to manufacturer’s instructions, with Seahorse XF RPMI, pH 7.4 supplemented with 1 mM pyruvate, 2 mM glutamine, and 10 mM glucose. GlycoPER was calculated using the Glycolytic Rate Assay report generator. All reagents were from Agilent.

### RNA isolation and RT-PCR

For RNA isolation, cells were washed with D-PBS and collected in TRI Reagent (Molecular Research Center). After addition of chloroform, total RNA was isolated from the aqueous phase using Econo-spin columns (Epoch Life Science) and subjected to on-column PureLink DNase (Invitrogen) digestion. To generate cDNA, 1 μg total RNA was reverse transcribed with MultiScribe Reverse Transcriptase (Applied Biosystems) with random hexamers (Invitrogen) and RNaseOUT (Invitrogen). Real-time PCR was performed using SYBR Green (Applied Biosystems) and the primers listed in table S1 on a ViiA 7 Real-Time PCR System (Applied Biosystems) with the following cycling parameters: 50°C (2 min), 95°C (10 min), 40 cycles of 95°C (15 sec) and 60°C (1 min), then dissociation curve analysis. Data were analyzed on QuantiStudio Real-Time PCR Software v1.3 (Applied Biosystems) and analyzed using the delta-delta Ct method. Isoforms of *XBP1* were detected using non-quantitative RT-PCR using the primers listed in table S1 and Phusion High-Fidelity PCR Master Mix (ThermoFisher) with the following cycling conditions: 98°C for 30 s, 35 cycles of 98°C (10 s), 65°C (30 s), and 72°C (30 s), then 72°C for 10 min. The resulting amplicons were separated and visualized on a 2.5%agarose gel containing SYBR Safe (Invitrogen).

### Protein isolation and western blots

For the TLR4 KO BMDMs, proteins were extracted from samples collected in TRI Reagent (Molecular Research Center) according to a modified protocol (65). For validation of the XBP1 KO and o/e lines, proteins were extracted using radioimmunoprecipitation assay (RIPA) buffer (50 mM Tris, 150 mM NaCl, 0.1% SDS, 0.5% sodium deoxycholate, and 1% Triton X-100) with Protease Inhibitor Cocktail Set III, Animal-Free (EMD Millipore). Insoluble debris was removed by centrifugation. Protein concentrations were determined using Pierce MicroBCA Protein Assay Kit (ThermoFisher). Equivalent amounts of protein were separated by SDS-PAGE and transferred to Immobilon-P PVDF membrane (Millipore). Membranes were incubated with antibodies per manufacturer’s suggestions. Blots were developed with Western Lightning Plus ECL (Perkin Elmer). The following antibodies were used: XBP1s (Cell Signaling Technology D2C1F; 12782), IRE1α (Cell Signaling Technology 14C10; 3294), GAPDH (Cell Signaling Technology 14C10; 2118), and goat anti-rabbit HRP (Jackson ImmunoResearch). Images were processed with Adobe Photoshop, which was utilized on occasion to change the order of lanes in the image to group appropriate samples together.

### Data analysis

Data was analyzed with Microsoft Excel (Microsoft) and Prism (GraphPad) with statistical tests indicated in figure legends. Densitometry was measured with ImageJ (version 1.53, ref. (66)). Data presented here are from a minimum of triplicate measurements from representative experiments.

## ACKNOWLEDGEMENTS

This work was supported by NIH grant AI109799 to RMT. HPS was supported by the National Center for Advancing Translational Sciences, National Institutes of Health, through grant number UL1 TR000002 and linked award TL1 TR000133. HPS was also supported by AI161850-01. BHA and MXO were supported by 1R21AI135403-01A1. Work in AJB’s laboratory was supported by award 650976 from the Crohn’s and Colitis Foundation of America NIH awards AI044170, AI096528, AI112445, AI112949 and AI146432. GS and JSK were supported by the Lupus Research Alliance. The content is solely the responsibility of the authors and does not necessarily represent the official views of the NIH.

We thank Dr. Bennett Penn and Andrew Rogers for sharing equipment, reagents, and ideas. We thank Sri Yalavarthi for help in acquiring reagents. We thank Dr. Xavier De Bolle for sharing the pKSoriT-*bla-kan-PsojA-mCherry* plasmid. We thank Bridget McLaughlin and the UC Davis flow cytometry core, which is funded by the UC Davis Comprehensive Cancer Center Support Grant (CCSG) awarded by the National Cancer Institute (NCI P30CA093373). We thank Dr. Alexey Tomilov and Dr. Gino Cortopassi for their help with the Seahorse assays. We thank Dr. Samantha Bell for guidance on representing CRISPR/Cas9-introduced mutations (67).

## REFERENCES

1. Viola A, Munari F, Sanchez-Rodriguez R, Scolaro T, Castegna A. 2019. The Metabolic Signature of Macrophage Responses. Front Immunol 10:1462.

2. Czyz DM, Willett JW, Crosson S. 2017. Brucella abortus Induces a Warburg Shift in Host Metabolism That Is Linked to Enhanced Intracellular Survival of the Pathogen. J Bacteriol 199.

3. Gomes MTR, Guimaraes ES, Marinho FV, Macedo I, Aguiar E, Barber GN, Moraes-Vieira PMM, Alves-Filho JC, Oliveira SC. 2021. STING regulates metabolic reprogramming in macrophages via HIF-1alpha during Brucella infection. PLoS Pathog 17:e1009597.

4. Xavier MN, Winter MG, Spees AM, den Hartigh AB, Nguyen K, Roux CM, Silva TM, Atluri VL, Kerrinnes T, Keestra AM, Monack DM, Luciw PA, Eigenheer RA, Baumler AJ, Santos RL, Tsolis RM. 2013. PPARgamma-mediated increase in glucose availability sustains chronic Brucella abortus infection in alternatively activated macrophages. Cell Host Microbe 14:159–70.

5. Adams CJ, Kopp MC, Larburu N, Nowak PR, Ali MMU. 2019. Structure and Molecular Mechanism of ER Stress Signaling by the Unfolded Protein Response Signal Activator IRE1. Front Mol Biosci 6:11.

6. Yoshida H, Matsui T, Yamamoto A, Okada T, Mori K. 2001. XBP1 mRNA Is Induced by ATF6 and Spliced by IRE1 in Response to ER Stress to Produce a Highly Active Transcription Factor. Cell 107:881–891.

7. Cox JS, Walter P. 1996. A Novel Mechanism for Regulating Activity of a Transcription Factor That Controls the Unfolded Protein Response. Cell 87:391–404.

8. Urano F, Wang X, Bertolotti A, Zhang Y, Chung P, Harding HP, Ron D. 2000. Coupling of stress in the ER to activation of JNK protein kinases by transmembrane protein kinase IRE1. Science 287:664–6.

9. Hu P, Han Z, Couvillon AD, Kaufman RJ, Exton JH. 2006. Autocrine tumor necrosis factor alpha links endoplasmic reticulum stress to the membrane death receptor pathway through IRE1alpha-mediated NF-kappaB activation and down-regulation of TRAF2 expression. Mol Cell Biol 26:3071–84.

10. Hu R, Warri A, Jin L, Zwart A, Riggins RB, Fang HB, Clarke R. 2015. NF-kappaB signaling is required for XBP1 (unspliced and spliced)-mediated effects on antiestrogen responsiveness and cell fate decisions in breast cancer. Mol Cell Biol 35:379–90.

11. Keestra-Gounder AM, Byndloss MX, Seyffert N, Young BM, Chavez-Arroyo A, Tsai AY, Cevallos SA, Winter MG, Pham OH, Tiffany CR, de Jong MF, Kerrinnes T, Ravindran R, Luciw PA, McSorley SJ, Baumler AJ, Tsolis RM. 2016. NOD1 and NOD2 signalling links ER stress with inflammation. Nature 532:394–7.

12. Pei G, Zyla J, He L, Moura-Alves P, Steinle H, Saikali P, Lozza L, Nieuwenhuizen N, Weiner J, Mollenkopf HJ, Ellwanger K, Arnold C, Duan M, Dagil Y, Pashenkov M, Boneca IG, Kufer TA, Dorhoi A, Kaufmann SH. 2021. Cellular stress promotes NOD1/2-dependent inflammation via the endogenous metabolite sphingosine-1-phosphate. EMBO J 40:e106272.

13. Martinon F, Chen X, Lee AH, Glimcher LH. 2010. TLR activation of the transcription factor XBP1 regulates innate immune responses in macrophages. Nat Immunol 11:411–8.

14. Zeng L, Liu YP, Sha H, Chen H, Qi L, Smith JA. 2010. XBP-1 couples endoplasmic reticulum stress to augmented IFN-beta induction via a cis-acting enhancer in macrophages. J Immunol 185:2324–30.

15. Song M, Sandoval TA, Chae CS, Chopra S, Tan C, Rutkowski MR, Raundhal M, Chaurio RA, Payne KK, Konrad C, Bettigole SE, Shin HR, Crowley MJP, Cerliani JP, Kossenkov AV, Motorykin I, Zhang S, Manfredi G, Zamarin D, Holcomb K, Rodriguez PC, Rabinovich GA, Conejo-Garcia JR, Glimcher LH, Cubillos-Ruiz JR. 2018. IRE1alpha-XBP1 controls T cell function in ovarian cancer by regulating mitochondrial activity. Nature 562:423–428.

16. Dong H, Adams NM, Xu Y, Cao J, Allan DSJ, Carlyle JR, Chen X, Sun JC, Glimcher LH. 2019. The IRE1 endoplasmic reticulum stress sensor activates natural killer cell immunity in part by regulating c-Myc. Nat Immunol 20:865–878.

17. Yan D, Wang HW, Bowman RL, Joyce JA. 2016. STAT3 and STAT6 Signaling Pathways Synergize to Promote Cathepsin Secretion from Macrophages via IRE1alpha Activation. Cell Rep 16:2914–2927.

18. Shan B, Wang X, Wu Y, Xu C, Xia Z, Dai J, Shao M, Zhao F, He S, Yang L, Zhang M, Nan F, Li J, Liu J, Liu J, Jia W, Qiu Y, Song B, Han JJ, Rui L, Duan SZ, Liu Y. 2017. The metabolic ER stress sensor IRE1alpha suppresses alternative activation of macrophages and impairs energy expenditure in obesity. Nat Immunol 18:519–529.

19. Batista A, Rodvold JJ, Xian S, Searles SC, Lew A, Iwawaki T, Almanza G, Waller TC, Lin J, Jepsen K, Carter H, Zanetti M. 2020. IRE1alpha regulates macrophage polarization, PD-L1 expression, and tumor survival. PLoS Biol 18:e3000687.

20. Choi JA, Song CH. 2019. Insights Into the Role of Endoplasmic Reticulum Stress in Infectious Diseases. Front Immunol 10:3147.

21. Celli J, Tsolis RM. 2015. Bacteria, the endoplasmic reticulum and the unfolded protein response: friends or foes? Nat Rev Microbiol 13:71–82.

22. Celli J. 2019. The Intracellular Life Cycle of Brucella spp. Microbiol Spectr 7.

23. Taguchi Y, Imaoka K, Kataoka M, Uda A, Nakatsu D, Horii-Okazaki S, Kunishige R, Kano F, Murata M. 2015. Yip1A, a novel host factor for the activation of the IRE1 pathway of the unfolded protein response during Brucella infection. PLoS Pathog 11:e1004747.

24. Li P, Tian M, Bao Y, Hu H, Liu J, Yin Y, Ding C, Wang S, Yu S. 2017. Brucella Rough Mutant Induce Macrophage Death via Activating IRE1alpha Pathway of Endoplasmic Reticulum Stress by Enhanced T4SS Secretion. Front Cell Infect Microbiol 7:422.

25. Smith JA, Khan M, Magnani DD, Harms JS, Durward M, Radhakrishnan GK, Liu YP, Splitter GA. 2013. Brucella induces an unfolded protein response via TcpB that supports intracellular replication in macrophages. PLoS Pathog 9:e1003785.

26. Guimaraes ES, Gomes MTR, Campos PC, Mansur DS, Dos Santos AA, Harms J, Splitter G, Smith JA, Barber GN, Oliveira SC. 2019. Brucella abortus Cyclic Dinucleotides Trigger STING-Dependent Unfolded Protein Response That Favors Bacterial Replication. J Immunol 202:2671–2681.

27. de Jong MF, Starr T, Winter MG, den Hartigh AB, Child R, Knodler LA, van Dijl JM, Celli J, Tsolis RM. 2013. Sensing of bacterial type IV secretion via the unfolded protein response. MBio 4:e00418–12.

28. Wells KM, He K, Pandey A, Cabello A, Zhang D, Yang J, Gomez G, Liu Y, Chang H, Li X, Zhang H, Feng X, da Costa LF, Metz R, Johnson CD, Martin CL, Skrobarczyk J, Berghman LR, Patrick KL, Leibowitz J, Ficht A, Sze SH, Song J, Qian X, Qin QM, Ficht TA, de Figueiredo P. 2022. Brucella activates the host RIDD pathway to subvert BLOS1-directed immune defense. Elife 11.

29. Abuaita BH, Schultz TL, O’Riordan MX. 2018. Mitochondria-Derived Vesicles Deliver Antimicrobial Reactive Oxygen Species to Control Phagosome-Localized Staphylococcus aureus. Cell Host Microbe 24:625-636 e5.

30. O’Neill LAJ, Artyomov MN. 2019. Itaconate: the poster child of metabolic reprogramming in macrophage function. Nat Rev Immunol 19:273–281.

31. Liu N, Li Y, Dong C, Xu X, Wei P, Sun W, Peng Q. 2016. Inositol-Requiring Enzyme 1-Dependent Activation of AMPK Promotes Brucella abortus Intracellular Growth. J Bacteriol 198:986–93.

32. Pandey A, Lin F, Cabello AL, da Costa LF, Feng X, Feng HQ, Zhang MZ, Iwawaki T, Rice-Ficht A, Ficht TA, de Figueiredo P, Qin QM. 2018. Activation of Host IRE1alpha-Dependent Signaling Axis Contributes the Intracellular Parasitism of Brucella melitensis. Front Cell Infect Microbiol 8:103.

33. Qin QM, Pei J, Ancona V, Shaw BD, Ficht TA, de Figueiredo P. 2008. RNAi screen of endoplasmic reticulum-associated host factors reveals a role for IRE1alpha in supporting Brucella replication. PLoS Pathog 4:e1000110.

34. Qiu Q, Zheng Z, Chang L, Zhao YS, Tan C, Dandekar A, Zhang Z, Lin Z, Gui M, Li X, Zhang T, Kong Q, Li H, Chen S, Chen A, Kaufman RJ, Yang WL, Lin HK, Zhang D, Perlman H, Thorp E, Zhang K, Fang D. 2013. Toll-like receptor-mediated IRE1alpha activation as a therapeutic target for inflammatory arthritis. EMBO J 32:2477–90.

35. Sule G, Abuaita BH, Steffes PA, Fernandes AT, Estes SK, Dobry C, Pandian D, Gudjonsson JE, Kahlenberg JM, O’Riordan MX, Knight JS. 2021. Endoplasmic reticulum stress sensor IRE1alpha propels neutrophil hyperactivity in lupus. J Clin Invest 131.

36. Cross BC, Bond PJ, Sadowski PG, Jha BK, Zak J, Goodman JM, Silverman RH, Neubert TA, Baxendale IR, Ron D, Harding HP. 2012. The molecular basis for selective inhibition of unconventional mRNA splicing by an IRE1-binding small molecule. Proc Natl Acad Sci U S A 109:E869–78.

37. Chen X, Iliopoulos D, Zhang Q, Tang Q, Greenblatt MB, Hatziapostolou M, Lim E, Tam WL, Ni M, Chen Y, Mai J, Shen H, Hu DZ, Adoro S, Hu B, Song M, Tan C, Landis MD, Ferrari M, Shin SJ, Brown M, Chang JC, Liu XS, Glimcher LH. 2014. XBP1 promotes triple-negative breast cancer by controlling the HIF1alpha pathway. Nature 508:103–107.

38. Zou C, Wang Y, Shen Z. 2005. 2-NBDG as a fluorescent indicator for direct glucose uptake measurement. J Biochem Biophys Methods 64:207–15.

39. Goldstein J, Hoffman T, Frasch C, Lizzio EF, Beining PR, Hochstein D, Lee YL, Angus RD, Golding B. 1992. Lipopolysaccharide (LPS) from Brucella abortus is less toxic than that from Escherichia coli, suggesting the possible use of B. abortus or LPS from B. abortus as a carrier in vaccines. Infect Immun 60:1385–9.

40. Moreno E, Berman DT, Boettcher LA. 1981. Biological activities of Brucella abortus lipopolysaccharides. Infect Immun 31:362–70.

41. Jakka P, Bhargavi B, Namani S, Murugan S, Splitter G, Radhakrishnan G. 2018. Cytoplasmic Linker Protein CLIP170 Negatively Regulates TLR4 Signaling by Targeting the TLR Adaptor Protein TIRAP. J Immunol 200:704–714.

42. Jakka P, Namani S, Murugan S, Rai N, Radhakrishnan G. 2017. The Brucella effector protein TcpB induces degradation of inflammatory caspases and thereby subverts non-canonical inflammasome activation in macrophages. J Biol Chem 292:20613–20627.

43. Lee JJ, Kim DH, Kim DG, Lee HJ, Min W, Rhee MH, Cho JY, Watarai M, Kim S. 2013. Toll-like receptor 4-linked Janus kinase 2 signaling contributes to internalization of Brucella abortus by macrophages. Infect Immun 81:2448–58.

44. Campos MA, Rosinha GM, Almeida IC, Salgueiro XS, Jarvis BW, Splitter GA, Qureshi N, Bruna-Romero O, Gazzinelli RT, Oliveira SC. 2004. Role of Toll-like receptor 4 in induction of cell-mediated immunity and resistance to Brucella abortus infection in mice. Infect Immun 72:176–86.

45. Salcedo SP, Marchesini MI, Degos C, Terwagne M, Von Bargen K, Lepidi H, Herrmann CK, Santos Lacerda TL, Imbert PR, Pierre P, Alexopoulou L, Letesson JJ, Comerci DJ, Gorvel JP. 2013. BtpB, a novel Brucella TIR-containing effector protein with immune modulatory functions. Front Cell Infect Microbiol 3:28.

46. Conde-Alvarez R, Arce-Gorvel V, Iriarte M, Mancek-Keber M, Barquero-Calvo E, Palacios-Chaves L, Chacon-Diaz C, Chaves-Olarte E, Martirosyan A, von Bargen K, Grillo MJ, Jerala R, Brandenburg K, Llobet E, Bengoechea JA, Moreno E, Moriyon I, Gorvel JP. 2012. The lipopolysaccharide core of Brucella abortus acts as a shield against innate immunity recognition. PLoS Pathog 8:e1002675.

47. Walter F, O’Brien A, Concannon CG, Dussmann H, Prehn JHM. 2018. ER stress signaling has an activating transcription factor 6alpha (ATF6)-dependent “off-switch”. J Biol Chem 293:18270–18284.

48. den Hartigh AB, Sun YH, Sondervan D, Heuvelmans N, Reinders MO, Ficht TA, Tsolis RM. 2004. Differential requirements for VirB1 and VirB2 during Brucella abortus infection. Infect Immun 72:5143–9.

49. Hiyoshi H, English BC, Diaz-Ochoa VE, Wangdi T, Zhang LF, Sakaguchi M, Haneda T, Tsolis RM, Baumler AJ. 2021. Virulence factors perforate the pathogen-containing vacuole to signal efferocytosis. Cell Host Microbe doi:10.1016/j.chom.2021.12.001.

50. Blouin CC, Page EL, Soucy GM, Richard DE. 2004. Hypoxic gene activation by lipopolysaccharide in macrophages: implication of hypoxia-inducible factor 1alpha. Blood 103:1124–30.

51. Papa S, Choy PM, Bubici C. 2019. The ERK and JNK pathways in the regulation of metabolic reprogramming. Oncogene 38:2223–2240.

52. Almanza A, Mnich K, Blomme A, Robinson CM, Rodriguez-Blanco G, Kierszniowska S, McGrath EP, Le Gallo M, Pilalis E, Swinnen JV, Chatziioannou A, Chevet E, Gorman AM, Samali A. 2022. Regulated IRE1alpha-dependent decay (RIDD)-mediated reprograming of lipid metabolism in cancer. Nat Commun 13:2493.

53. Bronner DN, Abuaita BH, Chen X, Fitzgerald KA, Nunez G, He Y, Yin XM, O’Riordan MX. 2015. Endoplasmic Reticulum Stress Activates the Inflammasome via NLRP3- and Caspase-2-Driven Mitochondrial Damage. Immunity 43:451–62.

54. Gomes MT, Campos PC, Oliveira FS, Corsetti PP, Bortoluci KR, Cunha LD, Zamboni DS, Oliveira SC. 2013. Critical role of ASC inflammasomes and bacterial type IV secretion system in caspase-1 activation and host innate resistance to Brucella abortus infection. J Immunol 190:3629–38.

55. Lobet E, Willemart K, Ninane N, Demazy C, Sedzicki J, Lelubre C, De Bolle X, Renard P, Raes M, Dehio C, Letesson JJ, Arnould T. 2018. Mitochondrial fragmentation affects neither the sensitivity to TNFalpha-induced apoptosis of Brucella-infected cells nor the intracellular replication of the bacteria. Sci Rep 8:5173.

56. Demars A, Vitali A, Comein A, Carlier E, Azouz A, Goriely S, Smout J, Flamand V, Van Gysel M, Wouters J, Abendroth J, Edwards TE, Machelart A, Hoffmann E, Brodin P, De Bolle X, Muraille E. 2021. Aconitate decarboxylase 1 participates in the control of pulmonary Brucella infection in mice. PLoS Pathog 17:e1009887.

57. Rajeeve K, Vollmuth N, Janaki-Raman S, Wulff TF, Baluapuri A, Dejure FR, Huber C, Fink J, Schmalhofer M, Schmitz W, Sivadasan R, Eilers M, Wolf E, Eisenreich W, Schulze A, Seibel J, Rudel T. 2020. Reprogramming of host glutamine metabolism during Chlamydia trachomatis infection and its key role in peptidoglycan synthesis. Nat Microbiol 5:1390–1402.

58. Eisele NA, Ruby T, Jacobson A, Manzanillo PS, Cox JS, Lam L, Mukundan L, Chawla A, Monack DM. 2013. Salmonella require the fatty acid regulator PPARdelta for the establishment of a metabolic environment essential for long-term persistence. Cell Host Microbe 14:171–182.

59. Escoll P, Song OR, Viana F, Steiner B, Lagache T, Olivo-Marin JC, Impens F, Brodin P, Hilbi H, Buchrieser C. 2017. Legionella pneumophila Modulates Mitochondrial Dynamics to Trigger Metabolic Repurposing of Infected Macrophages. Cell Host Microbe 22:302-316 e7.

60. Huang S, Xing Y, Liu Y. 2019. Emerging roles for the ER stress sensor IRE1alpha in metabolic regulation and disease. J Biol Chem 294:18726–18741.

61. Copin R, Vitry MA, Hanot Mambres D, Machelart A, De Trez C, Vanderwinden JM, Magez S, Akira S, Ryffel B, Carlier Y, Letesson JJ, Muraille E. 2012. In situ microscopy analysis reveals local innate immune response developed around Brucella infected cells in resistant and susceptible mice. PLoS Pathog 8:e1002575.

62. Shalem O, Sanjana NE, Hartenian E, Shi X, Scott DA, Mikkelson T, Heckl D, Ebert BL, Root DE, Doench JG, Zhang F. 2014. Genome-scale CRISPR-Cas9 knockout screening in human cells. Science 343:84–87.

63. Brinkman EK, Chen T, Amendola M, van Steensel B. 2014. Easy quantitative assessment of genome editing by sequence trace decomposition. Nucleic Acids Res 42:e168.

64. Campeau E, Ruhl VE, Rodier F, Smith CL, Rahmberg BL, Fuss JO, Campisi J, Yaswen P, Cooper PK, Kaufman PD. 2009. A versatile viral system for expression and depletion of proteins in mammalian cells. PLoS One 4:e6529.

65. Kopec AM, Rivera PD, Lacagnina MJ, Hanamsagar R, Bilbo SD. 2017. Optimized solubilization of TRIzol-precipitated protein permits Western blotting analysis to maximize data available from brain tissue. J Neurosci Methods 280:64–76.

66. Schneider CA, Rasband WS, Eliceiri KW. 2012. NIH Image to ImageJ: 25 years of image analysis. Nat Methods 9:671–5.

67. Bell SL, Lopez KL, Cox JS, Patrick KL, Watson RO. 2021. Galectin-8 Senses Phagosomal Damage and Recruits Selective Autophagy Adapter TAX1BP1 To Control Mycobacterium tuberculosis Infection in Macrophages. mBio 12:e0187120.

68. Mills EL, Ryan DG, Prag HA, Dikovskaya D, Menon D, Zaslona Z, Jedrychowski MP, Costa ASH, Higgins M, Hams E, Szpyt J, Runtsch MC, King MS, McGouran JF, Fischer R, Kessler BM, McGettrick AF, Hughes MM, Carroll RG, Booty LM, Knatko EV, Meakin PJ, Ashford MLJ, Modis LK, Brunori G, Sevin DC, Fallon PG, Caldwell ST, Kunji ERS, Chouchani ET, Frezza C, Dinkova-Kostova AT, Hartley RC, Murphy MP, O’Neill LA. 2018. Itaconate is an anti-inflammatory metabolite that activates Nrf2 via alkylation of KEAP1. Nature 556:113–117.

69. English BC, Van Prooyen N, Ord T, Ord T, Sil A. 2017. The transcription factor CHOP, an effector of the integrated stress response, is required for host sensitivity to the fungal intracellular pathogen Histoplasma capsulatum. PLoS Pathog 13:e1006589.

