## Supplemental Files for "The IRE1α-XBP1 signaling axis promotes glycolytic reprogramming in response to inflammatory stimuli"

### SUPPLEMENTARY MATERIALS

**Figure S1**. IRE1$\alpha$ supports *B. abortus* intracellular replication. WT or IRE1$\alpha$ KO RAW 264.7 cells (**A**) and WT (LysM-Cre$^-$ *Ern1*$^{fl/fl}$) or IRE1$\alpha$ KO (LysM-Cre$^+$ *Ern1*$^{fl/fl}$) BMDMs (**B**) were infected with *B. abortus* and intracellular bacterial burden was enumerated by colony forming units (CFU) at the indicated time points. Data points are means of triplicate wells $\pm$ SD. * $P \leq 0.05$, ** $P \leq 0.01$, *** $P \leq 0.001$, Student's two-tailed t-test.

**Figure S2**. The RNase activity of IRE1$\alpha$ supports the glycolytic response to *B. abortus* infection or LPS stimulation in macrophages. BMDMs were infected with *B. abortus* for 48 h (**A**) or stimulated with *Salmonella* LPS for 24 h (**B**) and concurrently treated with 50 $\mu$M 4$\mu$8c. Relative expression of the indicated genes normalized to uninfected or unstimulated controls was assessed by RT-qPCR. Data are presented as means of triplicate wells $\pm$ SD. * $P \leq 0.05$, ** $P \leq 0.01$, Student's two-tailed t-test.

**Figure S3**. Generation of XBP1 KO cells. (**A**) Electropherograms from WT and XBP1 KO RAW 264.7 cells showing the nonsense mutation introduced by CRISPR/Cas9. (**B, C**) WT and KO cells were treated with 500 nM thapsigargin (Tg) for 18 h. XBP1s protein levels were assessed by western blot. (**C**) The relative expression of the XBP1s target *ERdj4* normalized to untreated controls was assessed in Tg-treated cells by RT-qPCR. Data are presented as means of triplicate wells $\pm$ SD. * $P \leq 0.05$, ** $P \leq 0.01$, Student's two-tailed t-test.

**Figure S4**. Comparison of mCherry *B. abortus* strains. (**A**) mCherry fluorescence was measured from the mChe-expressing WT, *virB2* mutant, and complemented *virB2* strains, and the fluorescence signal was normalized to viable organisms as measured by CFUs. No statistical difference (ns), one-way ANOVA. (**B, C**) RAW 264.7 cells were infected with indicated strains at an MOI of 100 for the T4SS-sufficient strains and an MOI of 2000 for the T4SS-deficient strains for 48 h. (**B**) (Left) Representative FACS plots showing the identification of mCherry-negative, mCherry-low, and mCherry-high populations of RAW 264.7 cells infected as indicated. RAW 264.7 cells infected with the non-fluorescent complemented strain is shown as an mCherry-negative control. (Right) mCherry MFI for the mCherry-high population of RAW 264.7 cells infected with the indicated *B. abortus* strains, ** $P \leq 0.01$, Student's two-tailed t-test. (**C**) Representative FACS plots showing the identification of mCherry-negative, mCherry-low, and mCherry-high populations of RAW 264.7 cells infected with WT or the complemented *virB2* strain. RAW 264.7 cells infected with the non-fluorescent WT strain is shown as an mCherry-negative control.

**Figure S5**. Reduced glycolytic induction does not impair *B. abortus* intracellular growth. Intracellular replication was enumerated by colony forming units (CFU) in RAW 264.7 cells (**A**) or BMDMs (**B**) of the indicated genotypes. Data points are means of triplicate wells $\pm$ SD.

**Figure S6**. Model of how IRE$\alpha$-XBP1 supports glycolysis in response to inflammatory stimuli. LPS from *S.* Typhimurium signals through TLR4 to activate IRE1$\alpha$, leading to the production of the transcription factor XBP1s, which supports glycolytic reprogramming. *B. abortus* uses its T4SS to activate the IRE1α-XBP1s signaling axis, which promotes an increase in glycolysis in infected macrophages. TLR4 also supports glycolysis but is not required for IRE1α activation during *B. abortus* infection. Dashed lines represent putative IRE1α-independent pathways of TLR4-mediated glycolytic reprogramming, and dotted lines represent putative XBP1s-independent pathways of IRE1α-mediated glycolytic reprogramming.

**Table S1**

| Name | Sequence (5' - 3') | Purpose | Source |
|---|---|---|---|
| SecE up | GGTCATAATTCCGGCTTCAA | validation of pKSoriT-bla-kan-PsojA-mCherry insertion site | this study |
| SecE down | ATAAGCTGGTCGGCAAGAAA | validation of pKSoriT-bla-kan-PsojA-mCherry insertion site | this study |
| mChe | AAGCGCATGAACTCCTTGAT | validation of pKSoriT-bla-kan-PsojA-mCherry insertion site | this study |
| ActB F | AGAGGGAAATCGTGCGTGAC | RT-qPCR | (1) |
| ActB R | CAATAGTGATGACCTGGCCGT | RT-qPCR | (1) |
| Glut1 F | GCTGTGCTTATGGGCTTCTC | RT-qPCR | (1) |
| Glut1 R | CACATACATGGGCACAAAGC | RT-qPCR | (1) |
| Pfkfb3 F | AGCTGCCCGGACAAAACAT | RT-qPCR | (1) |

| | | | |
|---|---|---|---|
| Pfkfb3 R | CTCGGCTTTAGTGCTTCTGGG | RT-qPCR | (1) |
| Irg1 F | GCAACATGATGCTCAAGTCTG | RT-qPCR | (2) |
| Irg1 R | TGCTCCTCCGAATGATACCA | RT-qPCR | (2) |
| IL6 F | GAGGATACCACTCCCAACAGACC | RT-qPCR | this study |
| IL6 R | AAGTGCATCATCGTTGTTCATACA | RT-qPCR | this study |
| XBP1 F | GGCCTTGTGGTTGAGAACCAGGAG | XBP1 splicing | (3) |
| XBP1 R | GAATGCCCAAAAGGATATCAGACTC | XBP1 splicing | (3) |
| mNTC sgRNA F | CACCGTCCTGCGCGATGACCGTCGG | NTC sgRNA | (4) |
| mNTC sgRNA R | AAACCCGACGGTCATCGCGCAGGAC | NTC sgRNA | (4) |
| XBP1 sgRNA F | CACCGCGGCCTTGTGGTTGAGAACC | XBP1 sgRNA | this study |
| XBP1 sgRNA R | AAACGGTTCTCAACCACAAGGCCGC | XBP1 sgRNA | this study |
| XBP1 seq F | GGCTGAAATCTGCGAGTAGTA | XBP1 locus amplification/sequencing | this study |

| | | | |
|---|---|---|---|
| XBP1 seq R | AGGAACATCTGCCTGTAATGG | XBP1 locus amplification/sequencing | this study |
| XBP1 Gibson F | CTTTAAAGGAACCAATTCAGTCGACGCCACCATGGTGGTGGTGGCAGCG | XBP1s amplification for Gibson assembly | this study |
| XBP1 Gibson R | GGTCTAGATATCTCGAGTGCGGCCGCTTAGACACTAATCAGCTGGGGG | XBP1s amplification for Gibson assembly | this study |

**TABLE S1 REFERENCES**

1. Xavier MN, Winter MG, Spees AM, den Hartigh AB, Nguyen K, Roux CM, Silva TM, Atluri VL, Kerrinnes T, Keestra AM, Monack DM, Luciw PA, Eigenheer RA, Baumler AJ, Santos RL, Tsolis RM. 2013. PPARgamma-mediated increase in glucose availability sustains chronic Brucella abortus infection in alternatively activated macrophages. Cell Host Microbe 14:159-70.

2. Mills EL, Ryan DG, Prag HA, Dikovskaya D, Menon D, Zaslona Z, Jedrychowski MP, Costa ASH, Higgins M, Hams E, Szpyt J, Runtsch MC, King MS, McGouran JF, Fischer R, Kessler BM, McGettrick AF, Hughes MM, Carroll RG, Booty LM, Knatko EV, Meakin PJ, Ashford MLJ, Modis LK, Brunori G, Sevin DC, Fallon PG, Caldwell ST, Kunji ERS, Chouchani ET, Frezza C, Dinkova-Kostova AT, Hartley RC, Murphy MP, O'Neill LA. 2018. Itaconate is an anti-inflammatory metabolite that activates Nrf2 via alkylation of KEAP1. Nature 556:113-117.

3. English BC, Van Prooyen N, Ord T, Ord T, Sil A. 2017. The transcription factor CHOP, an effector of the integrated stress response, is required for host sensitivity to the fungal intracellular pathogen Histoplasma capsulatum. PLoS Pathog 13:e1006589.

4. Abuaita BH, Schultz TL, O'Riordan MX. 2018. Mitochondria-Derived Vesicles Deliver Antimicrobial Reactive Oxygen Species to Control Phagosome-Localized Staphylococcus aureus. Cell Host Microbe 24:625-636 e5.

Fig. S1

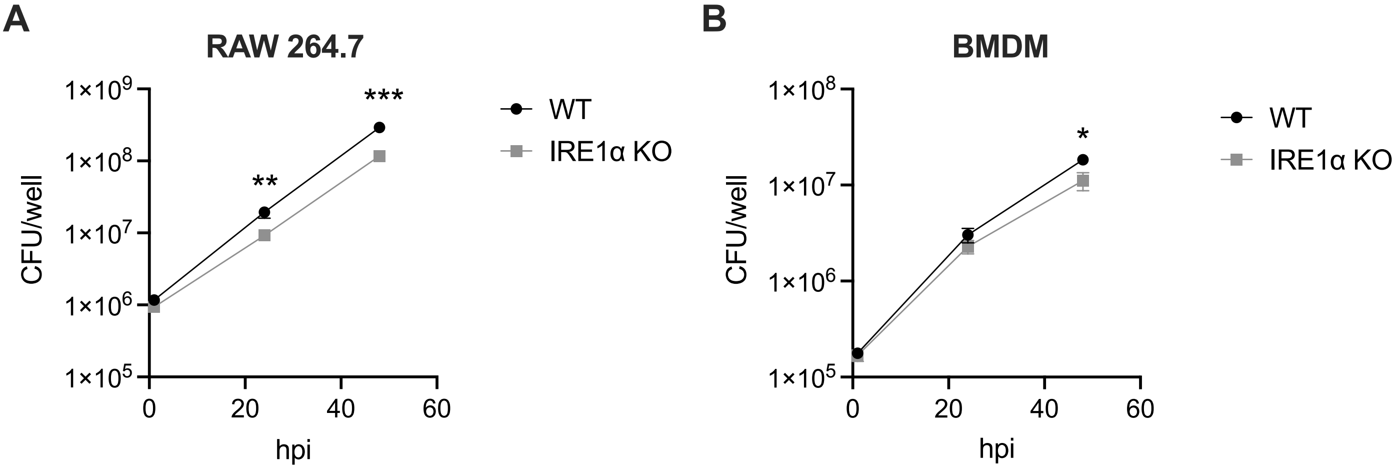
A
RAW 264.7
1×10^9
1×10^8
1×10^7
1×10^6
1×10^5
CFU/well
**
***
0
20
40
60
hpi
WT
IRE1α KO
B
BMDM
1×10^8
1×10^7
1×10^6
1×10^5
CFU/well
*
0
20
40
60
hpi
WT
IRE1α KO

Fig. S2

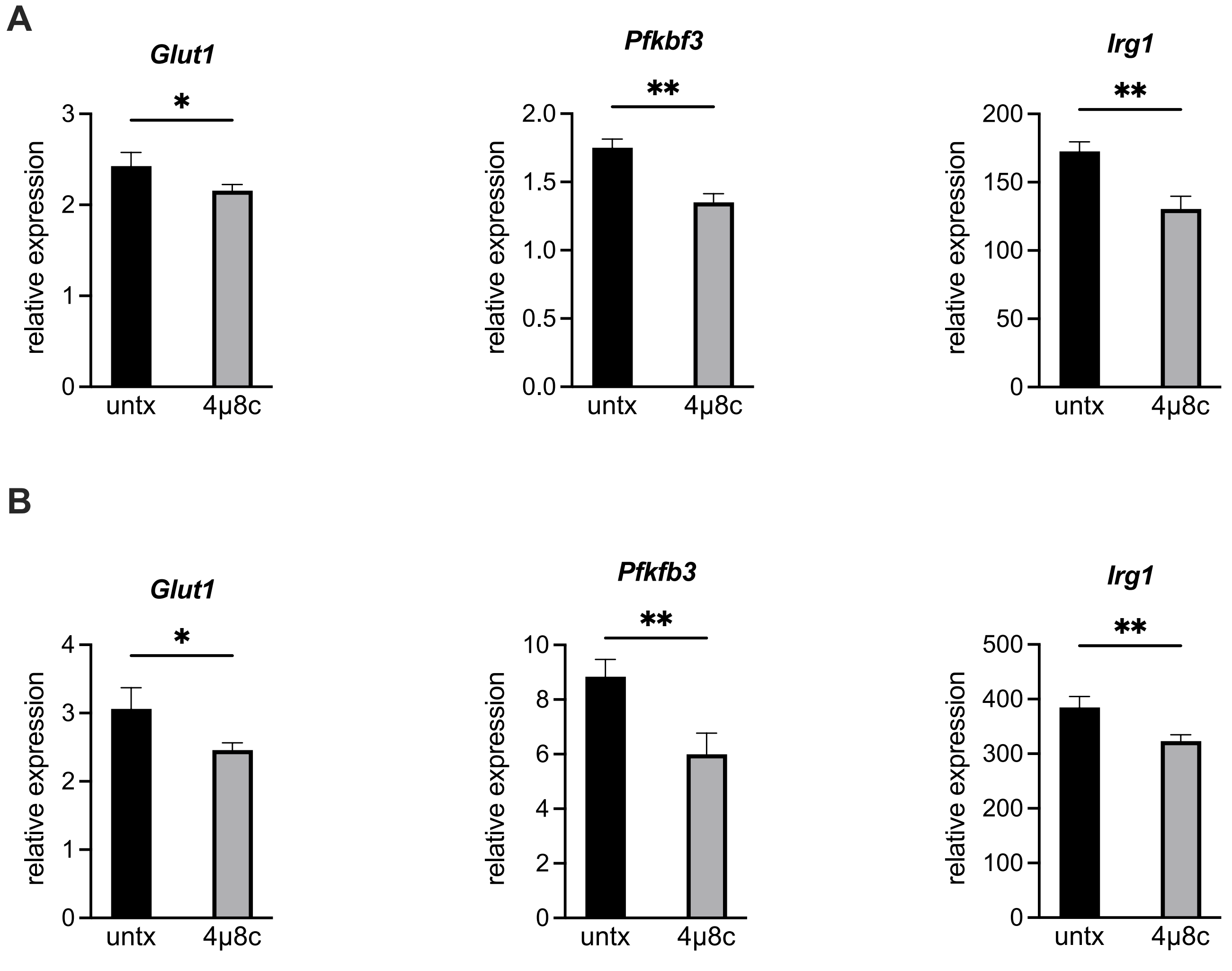
A
Glut1
*
relative expression
3
2
1
0
untx
4μ8c
Pfkbf3
**
relative expression
2.0
1.5
1.0
0.5
0.0
untx
4μ8c
Irg1
**
relative expression
200
150
100
50
0
untx
4μ8c
B
Glut1
*
relative expression
4
3
2
1
0
untx
4μ8c
Pfkfb3
**
relative expression
10
8
6
4
2
0
untx
4μ8c
Irg1
**
relative expression
500
400
300
200
100
0
untx
4μ8c

Fig. S3

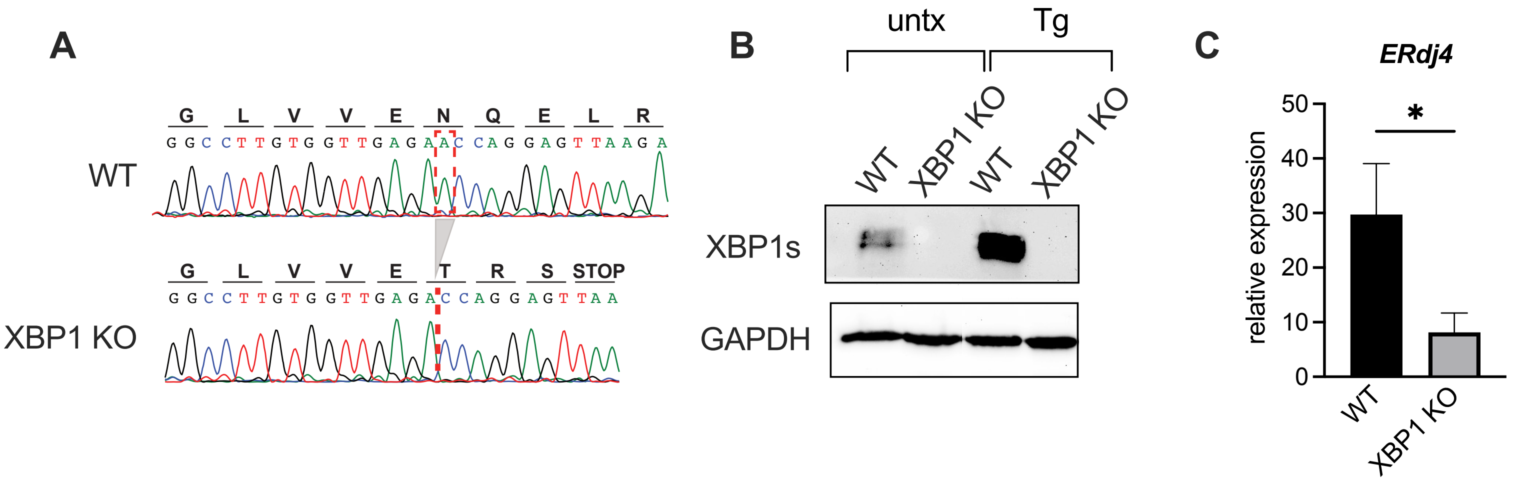
A
G L V V E N Q E L R
GGCCTTGTGGTTGAGAACCAGGAGTTAAGA
WT
G L V V E T R S STOP
GGCCTTGTGGTTGAGACCAGGAGTTAA
XBP1 KO
B
untx
Tg
WT
XBP1 KO
WT
XBP1 KO
XBP1s
GAPDH
C
ERdj4
relative expression
50
40
30
20
10
0
*
WT
XBP1 KO

Fig. S4

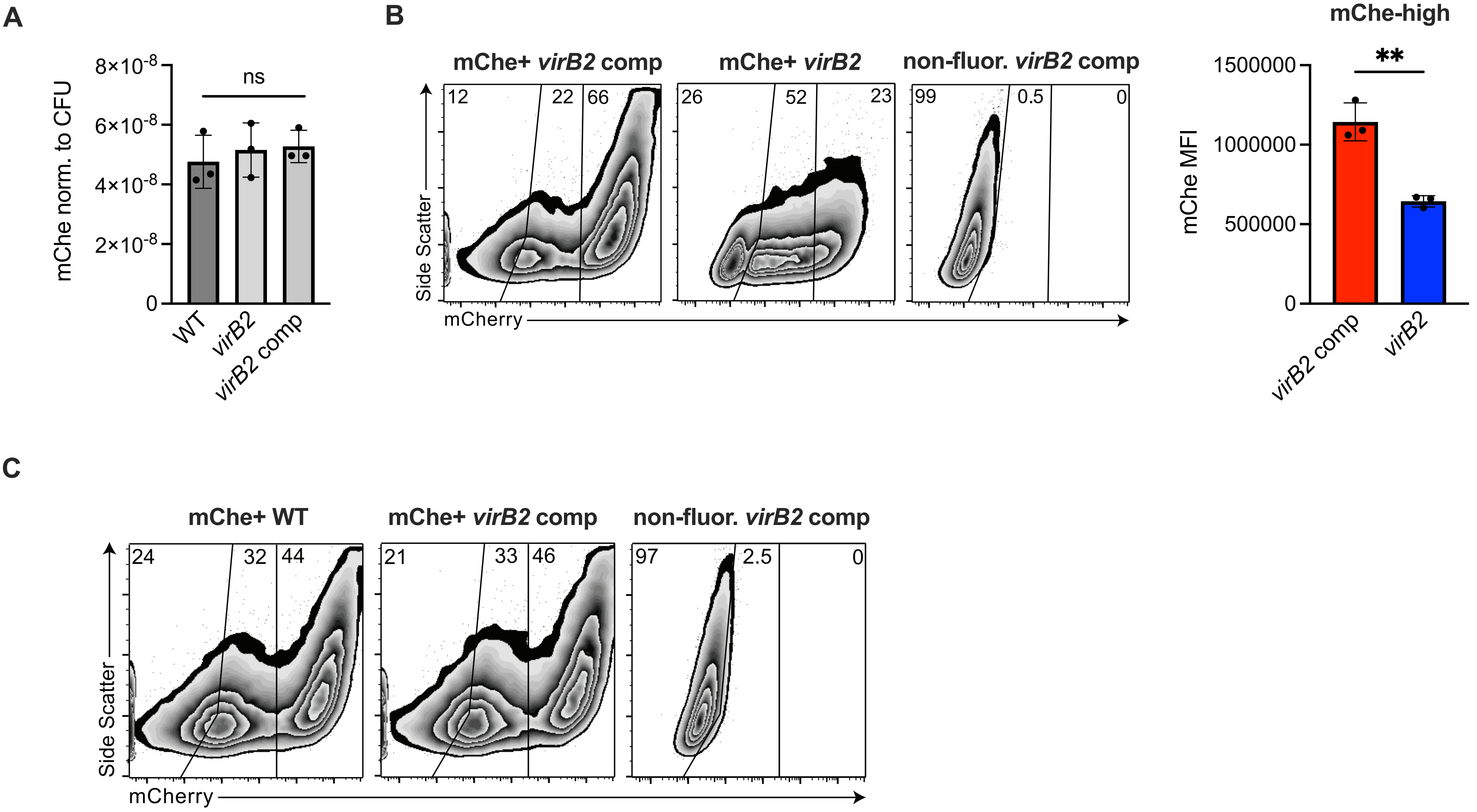
A
mChe norm. to CFU
8×10-8
6×10-8
4×10-8
2×10-8
0
ns
WT
virB2
virB2 comp
B
mChe+ virB2 comp
mChe+ virB2
non-fluor. virB2 comp
12
22
66
26
52
23
99
0.5
0
Side Scatter
mCherry
mChe-high
mChe MFI
1500000
1000000
500000
0
**
virB2 comp
virB2
C
mChe+ WT
mChe+ virB2 comp
non-fluor. virB2 comp
24
32
44
21
33
46
97
2.5
0
Side Scatter
mCherry

Fig. S5

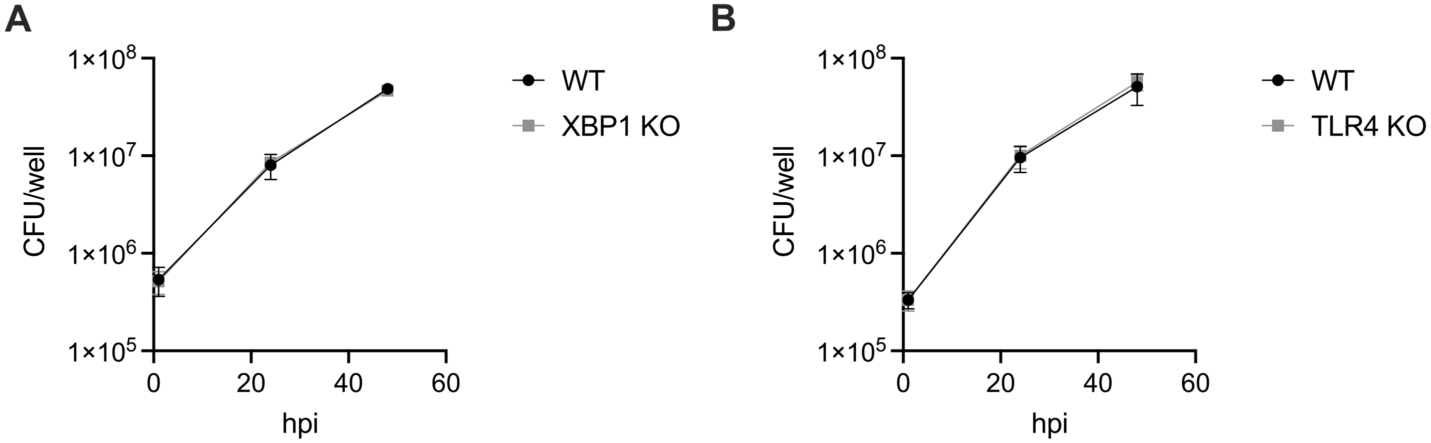
A
1×10^8
1×10^7
1×10^6
1×10^5
CFU/well
0
20
40
60
hpi
WT
XBP1 KO
B
1×10^8
1×10^7
1×10^6
1×10^5
CFU/well
0
20
40
60
hpi
WT
TLR4 KO

Fig. S6

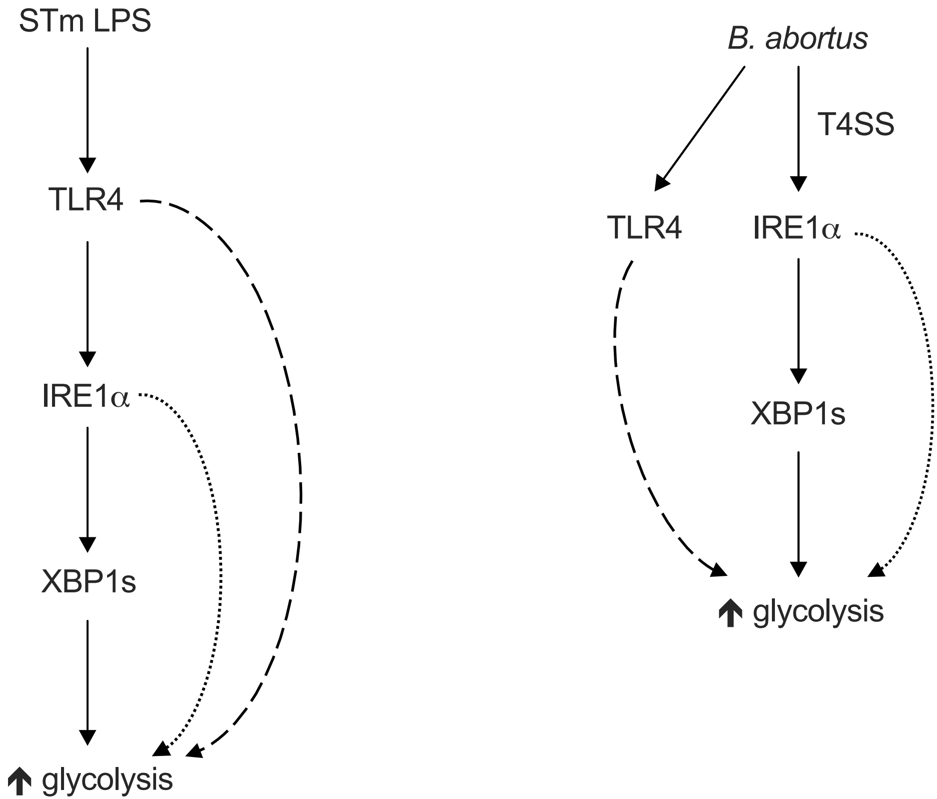
STm LPS
TLR4
IRE1α
XBP1s
glycolysis
B. abortus
T4SS
TLR4
IRE1α
XBP1s
glycolysis
